# Inositol pyrophosphate catabolism by three families of phosphatases controls plant growth and development

**DOI:** 10.1101/2024.04.30.591831

**Authors:** Florian Laurent, Simon M. Bartsch, Anuj Shukla, Felix Rico-Resendiz, Daniel Couto, Christelle Fuchs, Joël Nicolet, Sylvain Loubéry, Henning J. Jessen, Dorothea Fiedler, Michael Hothorn

**Author notes:** CSL Behring AG, Wankdorfstrasse 10, 3014, Bern, Switzerland.

## Abstract

Inositol pyrophosphates (PP-InsPs) are nutrient messengers whose cellular concentration must be tightly regulated. Diphosphoinositol pentakisphosphate kinases (PPIP5Ks) generate the active signaling molecule 1,5-InsP_8_. PPIP5Ks contain additional phosphatase domains involved in PP-InsP catabolism. Plant and Fungi Atypical Dual Specificity Phosphatases (PFA-DSPs) and NUDIX phosphatases (NUDTs) also hydrolyze PP-InsPs. Here we dissect the relative contributions of the three different phosphatase families to plant PP-InsP catabolism and nutrient signaling. We report the biochemical characterization of inositol pyrophosphate phosphatases from Arabidopsis and *Marchantia polymorpha*. Overexpression of different PFA-DSP and NUDT enzymes affects PP-InsP levels and leads to stunted growth phenotypes in Arabidopsis. *nudt17/18/21* knock-out mutants have altered PP-InsP pools and gene expression patterns, but no apparent growth defects. In contrast, *Marchantia polymorpha* Mp*pfa-dsp1^ge^,* Mp*nudt1^ge^* and Mp*vip1^ge^* mutants display severe growth and developmental phenotypes associated with changes in cellular PP-InsP levels. Analysis of Mp*pfa-dsp1^ge^*and Mp*vip1^ge^* supports a role for PP-InsPs in Marchantia phosphate signaling, and additional functions in nitrate homeostasis and cell wall biogenesis. Simultaneous removal of two phosphatase activities enhances the observed growth phenotypes. Taken together, PPIP5K, PFA-DSP and NUDT inositol pyrophosphate phosphatases play important roles in growth and development by collectively shaping plant PP-InsP pools.

**Author summary:** Organisms must maintain adequate levels of nutrients in their cells and tissues. One such nutrient is phosphorus, an essential building block of cell membranes, nucleic acids and energy metabolites. Plants take up phosphorus in the form of inorganic phosphate and require sufficient cellular phosphate levels to support their growth and development. It has been shown that plants and other eukaryotic organisms "measure" cellular phosphate levels using inositol pyrophosphate signaling molecules. The concentration of inositol pyrophosphates serves as a proxy for the cellular concentration of inorganic phosphate, and therefore inositol pyrophosphate synthesis and degradation must be tightly regulated. Here, we report that three different families of enzymes contribute to the degradation of inositol pyrophosphates in plants. The different phosphatases together shape cellular inositol pyrophosphate pools and thereby affect inorganic phosphate levels. Loss-of-function mutants of the different enzymes display additional defects in nitrate levels and cell wall architecture, suggesting that inositol pyrophosphates regulate cellular processes beyond inorganic phosphate homeostasis.

## Introduction

Inositol pyrophosphates (PP-InsPs) are small molecule nutrient messengers consisting of a fully phosphorylated *myo*-inositol ring and either one or two pyrophosphate groups(Shears, 2018). PP-InsPs are ubiquitous in eukaryotes where they perform diverse signaling functions. Their central role in cellular inorganic phosphate (Pi) / polyphosphate (polyP) homeostasis is conserved among fungi (Azevedo and Saiardi, 2017; Chabert et al., 2023; Guan et al., 2023; Wild et al., 2016), protozoa (Cordeiro et al., 2017), algae (Couso et al., 2016), plants (Stevenson-Paulik et al., 2005; Zhu et al., 2019; Dong et al., 2019; Ried et al., 2021; Guan et al., 2022) and animals (Gu et al., 2017; Haykir et al., 2024; Li et al., 2020; Wang et al., 2020).

In plants grown under Pi-sufficient conditions, the PP-InsP isomer 1,5-InsP_8_ accumulates in cells and binds to SPX (Syg1 Pho81 XPR1) receptor proteins (Dong et al., 2019; Ried et al., 2021; Wild et al., 2016). The ligand-bound receptor undergoes conformational changes (Pipercevic et al., 2023; Wild et al., 2016), for example allowing for the interaction with a family of PHOSPHATE STARVATION RESPONSE (PHR) transcription factors (Rubio et al., 2001; Lv et al., 2014; Puga et al., 2014; Wang et al., 2014; Wild et al., 2016). The coiled-coil oligomerisation and Myb DNA binding domains wrap around the SPX receptor, preventing PHRs from interacting with their target promoters. Under Pi starvation conditions, 1,5-InsP_8_ levels decrease, SPX – PHR complexes dissociate and the released transcription factors can oligomerize, bind promoters and regulate Pi starvation-induced (PSI) gene expression (Bustos et al., 2010; Ried et al., 2021; Guan et al., 2022).

Since cellular Pi homeostasis and 1,5-InsP_8_ levels are mechanistically linked, understanding the regulation of PP-InsP biosynthesis and catabolism is of fundamental and of biotechnological importance. In plants, PP-InsPs are generated from phytic acid (InsP_6_) by a series of pyrophosphorylation steps catalyzed by inositol 1,3,4-trisphosphate 5/6-kinases (ITPKs) (Laha et al., 2019) and by the diphosphoinositol pentakisphosphate kinases (PPIP5K) VIH1/2 (or VIP1/2) (Desai et al., 2014; Laha et al., 2015; Zhu et al., 2019; Dong et al., 2019). Consistent with the function of 1,5-InsP_8_ as a nutrient messenger in Pi homeostasis and starvation responses, deletion of enzymes that disrupt the biosynthesis of InsP_6_ (IPK1 and IPK2β), 5-InsP_7_ (ITPK1) or 1,5-InsP_8_ (VIH1/VIH2), results in altered Pi starvation responses in Arabidopsis (Stevenson-Paulik et al., 2005; Kuo et al., 2014, 2018; Zhu et al., 2019; Riemer et al., 2021; Dong et al., 2019). *vih1 vih2* loss-of-function mutants lack 1,5-InsP_8_, display constitutive Pi starvation responses and a severe seedling lethal phenotype, which can be partially rescued upon additional deletion of *PHR1* and its paralog *PHL1* (Bustos et al., 2010; Zhu et al., 2019).

PP-InsP catabolic enzymes have been identified in the C-terminus of PPIP5Ks (Mulugu et al., 2007), and as stand-alone enzymes in the Plant & Fungi Atypical Dual Specificity Phosphatase (PFA-DSPs) (Steidle et al., 2016) and the NUDIX (NUcleoside DIphosphates associated to moiety-X) hydrolase (hereafter NUDT) (Ingram et al., 1999; Cartwright and McLennan, 1999; Safrany et al., 1999) families. The fission yeast PPIP5K Asp1 has been characterized as a inositol 1-pyrophosphate phosphatase, releasing 5-InsP_7_ from 1,5-InsP_8_ and InsP_6_ from 1-InsP_7_ (Pöhlmann et al., 2014; Dollins et al., 2020).

The fungal PFA-DSPs ScSiw14 and SpSiw14 are metal-independent cysteine-dependent phosphatases capable of hydrolyzing 1-InsP_7_, 5-InsP_7_ and 1,5-InsP_8_ with a preference for 5-InsP_7_ (Sanchez et al., 2023; Steidle et al., 2016; Wang et al., 2018). The preferred substrate of the five PFA-DSPs in Arabidopsis is 5-InsP_7_ in the presence of Mg^2+^ ions (Kurz et al., 2023) *in vitro* (Gaugler et al., 2022; Wang et al., 2022).

NUDIX hydrolases are a large family of enzymes that share a common fold and broad substrate specificity (Carreras-Puigvert et al., 2017; Yoshimura and Shigeoka, 2015). NUDT enzymes of the diadenosine and diphosphoinositol polyphosphate phosphohydrolase subfamily have been characterized as inositol pyrophosphate phosphatases: fungal Ddp1 (Cartwright and McLennan, 1999) (YOR162w) and Aps1 (Ingram et al., 1999; Safrany et al., 1999) are able to hydrolyze different polyphosphate substrates, such as polyP, diadenosine polyphosphates (Ap_n_A) and inositol pyrophosphates, with a moderate substrate preference for 1-InsP_7_ (Ingram et al., 1999; Safrany et al., 1999; Garza et al., 2009; Lonetti et al., 2011; Kilari et al., 2013; Márquez-Moñino et al., 2021; Zong et al., 2021). Of the 28 NUDIX enzymes present in Arabidopsis (Yoshimura and Shigeoka, 2015), AtNUDT13 has been characterized as an Ap_6_A phosphohydrolase (Olejnik et al., 2007).

PPIP5K, PFA-DSP and NUDT phosphatase mutants have been characterized in fungi and in plants. Mutation of catalytic histidine in the phosphatase domain of fission yeast PPIP5K Asp1 altered microtubule dynamics and vacuolar morphology (Pascual-Ortiz et al., 2018; Dollins et al., 2020). Severe growth phenotypes have been reported for missense alleles leading to early stop mutations in phosphatase domain of Asp1 (Garg et al., 2020). In Arabidopsis, complementation of the seedling lethal phenotype of *vih1-2 vih2-4* mutant plants with the full-length PPIP5K VIH2 containing a catalytically inactive phosphatase domain restored growth back to wild-type levels with only minor Pi accumulation defects (Zhu et al., 2019). Baker’s yeast PFA-DSP *siw14*Δ strains showed enhanced environmental stress responses (Steidle et al., 2020) and increased 5-InsP_7_ levels (Chabert et al., 2023). T-DNA insertion lines in the *pfa-dsp1* locus had no apparent phenotypes and wild-type-like cellular PP-InsP levels (Gaugler et al., 2022). Over-expression of *AtPFA-DSP1* in Arabidopsis or in *Nicotiana benthamiana* resulted in decreased InsP_7_ pools (Gaugler et al., 2022). Overexpression of *AtPFA-DSP4*, or of rice *OsPFA-DSP1* and *OsPFA-DSP2* resulted in altered drought and pathogen responses (He et al., 2012; Liu et al., 2012).

Genetic interaction studies between PPIP5Ks, PFA-DSPs and NUDIX enzymes have been performed in fungi. In baker’s yeast, *siw14Δ vip1Δ* and *siw14Δ ddp1Δ* contained higher cellular InsP_7_ levels when compared to the respective single mutants (Steidle et al., 2016). In fission yeast, neither the Asp1, Aps1 nor the Siw14 phosphatase activities were required for vegetative growth (Sanchez et al., 2023). Importantly, *aps1Δ asp1-H297A* double mutants are lethal and this phenotype is dependent on 1,5-InsP_8_ synthesis by the PPIP5K Asp1 (Sanchez et al., 2019). Likewise, *aps1*Δ *siw14-C189S* mutants are lethal, suggesting that combined 1-InsP_7_ and 5-InsP_7_ catabolism is essential in fission yeast (Sanchez et al., 2023).

The relative contributions of PPIP5K, PFA-DSP and NUDT inositol pyrophosphate phosphatases to plant PP-InsP catabolism remain to be characterized. Here, using a PP-InsP affinity reagent previously developed to identify PP-InsP interacting proteins in yeast (Wu et al., 2016) and in human cells (Furkert et al., 2020), we isolate three PFA-DSP and three NUDT inositol pyrophosphate phosphatases from Arabidopsis and characterize their *in vitro* enzyme properties and *in planta* gain- and loss-of-function phenotypes. Translating our findings to *Marchantia polymorpha*, we define loss-of-function phenotypes for PFA-DSP, NUDT and PPIP5K phosphatases and investigate their genetic interaction.

## Results

### AtPFA-DSP1 and AtNUDT17 are inositol pyrophosphate phosphatases

To identify putative inositol pyrophosphate phosphatases in Arabidopsis we prepared protein extracts from 2-week-old seedlings grown under Pi-sufficient or Pi starvation conditions and performed affinity pull-downs with resin-immobilized 5PCP-InsP_5_, a non-hydrolyzable PP-InsP analog (Wu et al., 2016) (Supplementary Figure 1A, see Methods). Different InsP/PP-InsP kinases including ITPK1/2 (Laha et al., 2019) and VIH1/2 (Laha et al., 2015; Desai et al., 2014; Zhu et al., 2019; Dong et al., 2019) specifically bound to 5PCP-InsP_5_ but not to Pi control beads (Supplementary Figure 1B). Six putative PP-InsP phosphatases were recovered, including AtPFA-DSP1, AtPFA-DSP2 and AtPFA-DSP4 as well as AtNUDT17, AtNUDT18 and AtNUDT21 (Supplementary Figure 1B). We excluded several purple acid phosphatases from further analysis (Supplementary Figure 1B), because they are likely cell wall-resident enzymes involved in Pi foraging (Del Vecchio et al., 2014). Samples from Pi-starved and Pi-sufficient conditions all contained the different PP-InsP metabolizing enzymes, but their protein abundance was overall increased under Pi starvation.

We next tested whether AtPFA-DSP1/2/4 and AtNUDT17/18/21 are inositol pyrophosphate phosphatases *in vitro* (Supplementary Figure 1C). Therefore, we expressed and purified recombinant AtPFA-DSP1 (residues 1-216) and AtNUDT17 (residues 23-163) and characterized their enzyme activities (see Methods, Supplementary Figure 2). We found that both AtPFA-DSP1 and AtNUDT17 are inositol pyrophosphate phosphatases with 5-InsP_7_ being the preferred substrate for both enzymes *in vitro* (see below, Figure 1A, B and Supplementary Figure 2). Both enzymes do not require a metal co-factor for catalysis (Lonetti et al., 2011; Steidle et al., 2016). However, the conformational equilibrium of PP-InsPs can be modulated by metal cations (Kurz et al., 2023) and hence we performed enzyme assays in the presence and absence of MgCl_2_ (Figure 1A, B). Taken together, AtPFA-DSP1 and AtNUDT17 are *bona fide* inositol pyrophosphate phosphatases.

**Figure 1.**
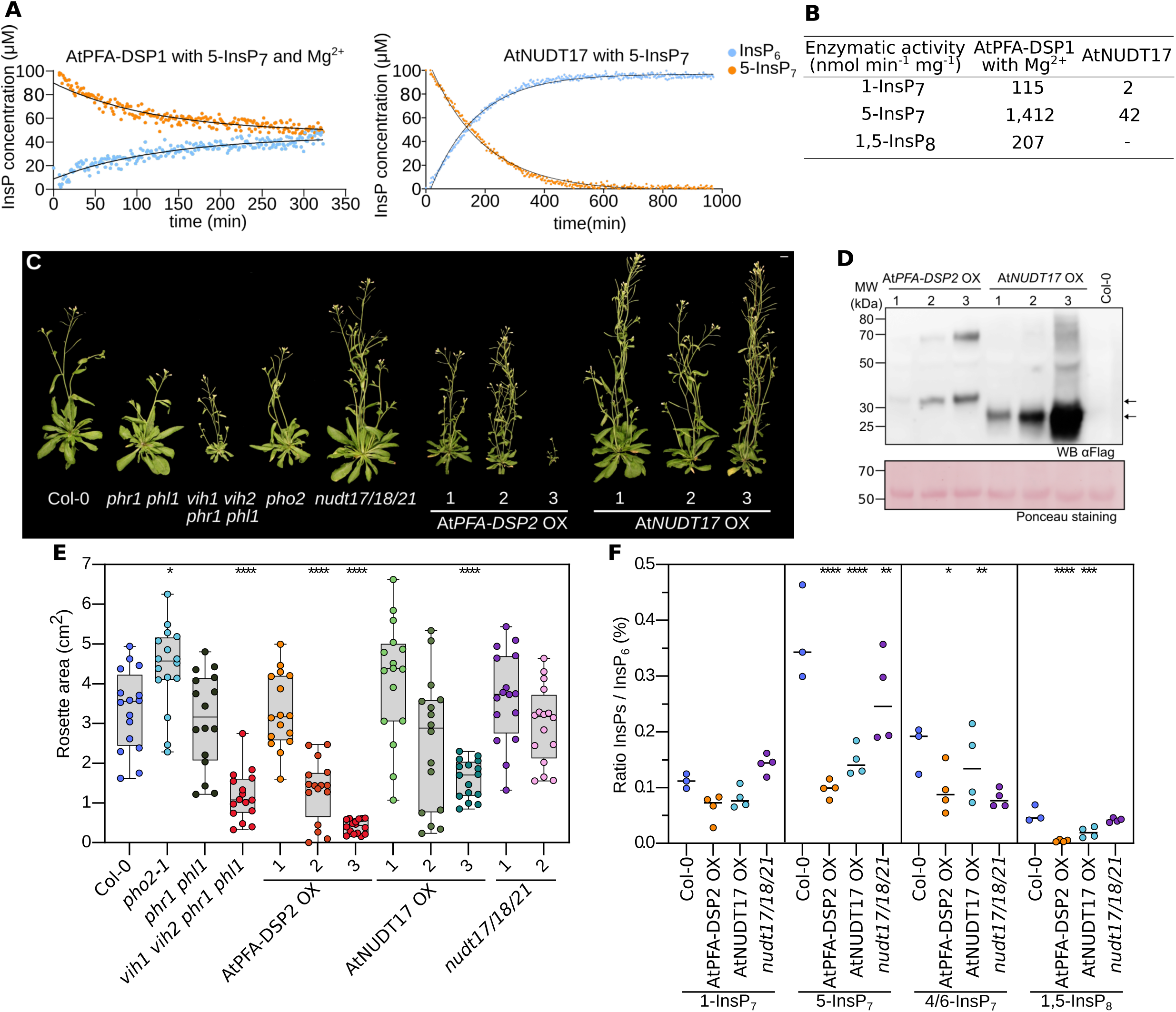
Overexpressing inositol pyrophosphate phosphatases restricts Arabidopsis growth and alters PP-InsP levels. **(A)** NMR-based inositol phosphatase assays. Shown are time course experiments of AtPFA-DSP1 and AtNUDT17 using 100 μM of [^13^C_6_] 5-InsP_7_ as substrate. Pseudo-2D spin-echo difference experiments were used and the relative intensity changes of the C2 peaks of InsP_6_ and 5-InsP_7_ as function of time were quantified. **(B)** Table summaries of the enzymatic activities of AtPFA-DSP1 and AtNUDT17 vs. PP-InsPs substrates. **(C)** Growth phenotypes of 4-week-old *nudt17/18/21*, At*PFA-DSP2* OX, At*NUDT17* OX plants. *phr1 phl1*, *vih1 vih2 phr1 phl1* and *pho2* mutants and Col-0 plants of the same age are shown as controls. Plants were germinated on ^½^MS for one week before transferring to soil for additional 3 weeks. Scale bar = 1 cm. **(D)** Western blot of At*PFA-DSP2* OX and At*NUDT17* OX plants vs. the Col-0 control. AtPFA-DSP2-Flag has a calculated molecular mass of ∼31 kDa and AtNUDT17-Flag ∼24 kDa. A Ponceau stain is shown as loading control below. Arrows indicate the expected sizes of AtPFA-DSP2 (top) and AtNUDT17 (bottom). **(E)** Rosette surface areas of 3-week-old *nudt17/18/21*, At*PFA-DSP2* OX and At*NUDT17* OX plants, controls as in **(C)** Multiple comparisons of the genotypes vs. wild-type (Col-0) were performed according Dunnett (Dunnett, 1955) test as implemented in the R package multcomp (Hothorn et al., 2008) (**** p < 0.001, *** p < 0.005, ** p < 0.01, * p < 0.05). **(F)** Whole tissue PP-InsP quantification of 2-week-old Col-0, *nudt17/18/21*, AtPFA-DSP2 OX and AtNUDT17 OX seedlings. PP-InsP levels were normalized by InsP_6_ levels.

### Overexpression of AtPFA-DSPs or AtNUDTs results in stunted growth and altered PP-InsP pools

Arabidopsis AtPFA-DSP1/2/4 and AtNUDT17/18/21 group with their respective Siw14 and Ddp1 orthologs from yeast in phylogenetic trees, respectively (Supplementary Figure 1D-G). We next used <u>c</u>lustered regularly inter<u>s</u>paced <u>p</u>alindromic repeats (CRISPR/Cas9) gene editing (Jinek et al., 2012) to generate *nudt17/18/21* triple loss-of-function mutants (Supplementary Figure 3), and *AtNUDT17*, *AtNUDT18* and *AtNUDT21* overexpression (OX) lines (Figure 1c, Supplementary Figure 4A, B). We also generated ubiquitin 10 promoter-driven *AtPFA-DSP1*, *AtPFA-DSP2* and *AtPFA-DSP4* OX lines, but were unable to isolate higher order *pfa-dsp1/2/4* mutants (Figure 1C, Supplementary Figure 4A, B). *nudt17/18/21* loss-of-function mutants and *AtNUDT17*, *AtNUDT18* or *AtNUDT21* OX lines showed no severe growth phenotypes (Figure 1A and Supplementary Figure 4A). Overexpression of either *AtPFA-DSP1*, *AtPFA-DSP2* or *AtPFA-DSP4* resulted in stunted growth phenotypes (Figure 1C and Supplementary Figure 4A, B). *AtPFA-DSP2* OX and *AtNUDT17* OX lines both exhibited reduced rosette areas, which positively correlated with the protein expression level in the respective independent T3 line (Figure 1D, E). Overexpression of AtPFA-DSP1 in Arabidopsis has previously been associated with a reduction in cellular InsP_7_ pools (Gaugler et al., 2022). We therefore quantified cellular PP-InsP levels by capillary electrophoresis coupled to mass spectrometry in our different transgenic lines (Qiu et al., 2023, 2020). *AtPFA-DSP2* OX lines showed reduced levels of 5-InsP_7_ and 1,5-InsP_8_, in good agreement with the inositol 5-pyrophosphate phosphatase activity of this enzyme *in vitro* (Figure 1A, B, F and Supplementary Figure 5). Consistent with our biochemical assays, *AtNUDT17* OX lines also showed reduced 5-InsP_7_ levels (Figure 1A, B, F and Supplementary Figure 5). Only minor changes in PP-InsP pools were observed in *nudt17/18/21* plants, but InsP_6_ levels were elevated (Figure 1F and Supplementary Figure 5). *AtNUDT17*, *AtNUDT18* and *AtNUDT21* are expressed at seedling stage as concluded from the analysis of promoter::β-glucuronidase (GUS) fusions (Figure 2A).

**Figure 2.**
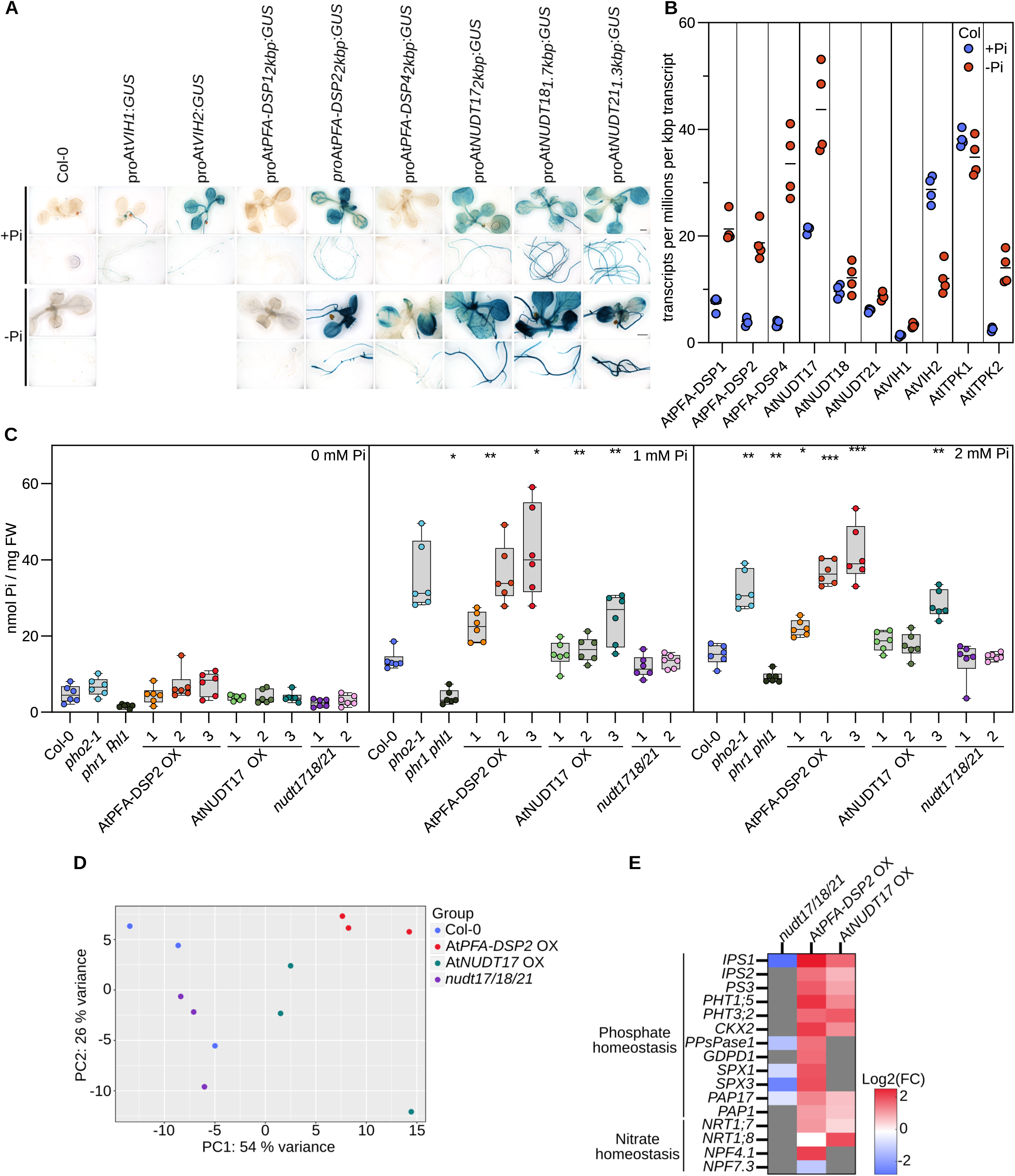
AtPFA-DSPs and AtNUDTs regulate Pi homeostasis in Arabidopsis. **(A)** Promoter β-glucuronidase (GUS) reporter assay for 2-week-old At*PFA-DSP1/2/4* OX and At*NUDT17/18/21* OX seedlings. The previously reported _pro_At*VIH1*::GUS and _pro_At*VIH2*::GUS lines (Zhu et al., 2019) are shown alongside. **(B)** Quantification of AtPFA-DSP1/2/4, AtNUDT17/18/21, AtVIH1/2 and AtITPK1/2 transcripts from RNA-seq experiments performed on 2-week-old Col-0 seedling grown in either no phosphate (-Pi) or in 1 mM K_2_HPO_4_/KH_2_PO_4_ (+Pi). Counts were normalized by the number of reads in each dataset and by the length of each transcript. **(C)** Total Pi concentrations of 2-week-old *nudt17/18/21*, At*PFA-DSP2* OX and At*NUDT17* OX seedlings grown in different Pi conditions. *phr1 phl1*, *vih1 vih2 phr1 phl1, pho2* and Col-0 plants were used as control. For each genotype and condition, 6 biological replicates from 3-4 pooled seedlings were used, technical triplicates were done for the standards and duplicates for all samples. A Dunnett test was performed to assess the statistical difference of the genotypes compared to Col-0 (**** p < 0.001, *** p < 0.005, ** p < 0.01, * p < 0.05). **(D)** Principal component analysis (PCA) of an RNA-seq experiment comparing 2-week-old *nudt1/18/21*, At*PFA-DSP2* OX and At*NUDT17* OX seedlings grown under Pi-sufficient conditions to the Col-0 reference. The read variance analysis was performed with DESeq2 and displayed with ggplot2 in R. **(E)** Heatmap of differentially expressed genes (DEGs) involved in Pi or Nitrogen homeostasis using the RNA-seq data from **(D)**. Known marker genes significantly different from Col-0 involved in Pi or Nitrogen homeostasis are displayed. Grey boxes = not differentially expressed from Col-0.

Our reporter lines showed that expression of all three NUDT genes as well as *AtPFA-DSP2* and *AtPFA-DSP4* is up-regulated under Pi starvation conditions (Figure 2A). Consistent with this, RNA-seq experiments comparing 2-week-old Col-0 seedlings grown in Pi sufficient vs. starved conditions showed increased transcript levels for *AtPFA-DSP1/2/4* and for *AtNUDT17* under Pi starvation (Figure 2B). We therefore quantified cellular Pi levels in our transgenic lines and found that similar to previously reported *vih1 vih2 (Zhu et al., 2019)* loss-of-function and constitutively active *PHR1 (Ried et al., 2021)* alleles, *AtPFA-DSP2* OX and *AtNUDT17* OX but not *nudt17/18/21* plants overaccumulate Pi in phosphate-sufficient growth conditions, when compared to the Col-0 control (Figure 2C). We hypothesized that Pi overaccumulation in *AtPFA-DSP2* OX and in *AtNUDT17* OX may be caused by reduced 1,5-InsP_8_ pools (Figure 1F), which in turn may lead to a constitutive activation of PHR1/PHL1 transcription factors (Guan et al., 2022; Ried et al., 2021; Wild et al., 2016; Zhu et al., 2019). We performed additional RNA-seq analyses and found that several conserved PSI marker genes such as *AtPPsPase1*, *AtSPX1*, *AtSPX3*, *AtIPS1* and *AtPHT1;5* were strongly up-regulated in *AtPFA-DSP2* OX and to a lesser extent in *AtNUDT17* OX lines (Figure 2D, E). Several PSI marker genes are repressed in the *nudt17/18/21* knock-out line (Figure 2E). Notably, we also observed induction of nitrate transporters in *AtPFA-DSP2* OX and in *AtNUDT17* OX plants (Figure 2E). Taken together, AtPFA-DSP1/2/4 or AtNUDT17/18/21 overexpression can alter PP-InsP pools and cellular responses.

### Identification of PFA-DSP and NUDT inositol pyrophosphate phosphatases in Marchantia

Characterization of our *nudt17/18/21* triple mutant revealed no obvious visual or molecular phenotypes (Figure 1), suggesting that other members of the large Arabidopsis NUDIX gene family (Yoshimura and Shigeoka, 2015) may have redundant inositol pyrophosphate phosphatase activities. Indeed, biochemical analysis of AtNUDT13 (residues 1-202), which was previously characterized as an Ap_6_A phosphohydrolase (Olejnik et al., 2007), revealed robust inositol 1- and 5-pyrophosphate phosphatase activity, that exceeded that observed for AtNUDT17 (Supplementary Figure 2B, C).

To overcome the potential genetic redundancies within the Arabidopsis PFA-DSP and NUDIX enzyme families, we sought to identify *bona fide* inositol pyrophosphate phosphatases in the liverwort *Marchantia polymorpha*. Using phylogenetic trees derived from multiple sequence alignments, we identified Mp3g10950 (https://marchantia.info, hereafter Mp*PFA-DSP1*) in the subtree containing the Arabidopsis PFA-DSPs and ScSiw14 (Supplementary Figure 1D). Similarly, Mp5g06600 (Mp*NUDT1*) clusters with *AtNUDT17*, *AtNUDT18*, *AtNUDT21* and with yeast Ddp1 (Supplementary Figure 1E). We expressed and purified recombinant *M. polymorpha* MpPFA-DSP1 (residues 4-171) and MpNUDT1 (18-169) and evaluated their inositol pyrophosphate phosphatase activities (Supplementary Figure 6). MpPFA-DSP1 is a specific inositol 5-pyrophosphate phosphatase with a substrate preference for 5-InsP_7_ over 1,5-InsP_8_ (Figure 3A, B and Supplementary Figure 6). Mutation of the catalytic Cys105 to Ala rendered MpPFA-DSP1 catalytically inactive (Figure 3b and Supplementary Figure 6). In contrast to AtNUDT17 or AtNUDT13 (Figure 1B and Supplementary Figure 2C, D), MpNUDT1 is an inositol 1-pyrophosphate phosphatase that cleaves 1-InsP_7_, an activity that depends on the catalytic Glu79 (Figure 3A, B and Supplementary Figure 6). In conclusion, MpPFA-DSP1 and MpNUDT1 are specific inositol pyrophosphate phosphatases in Marchantia.

**Figure 3.**
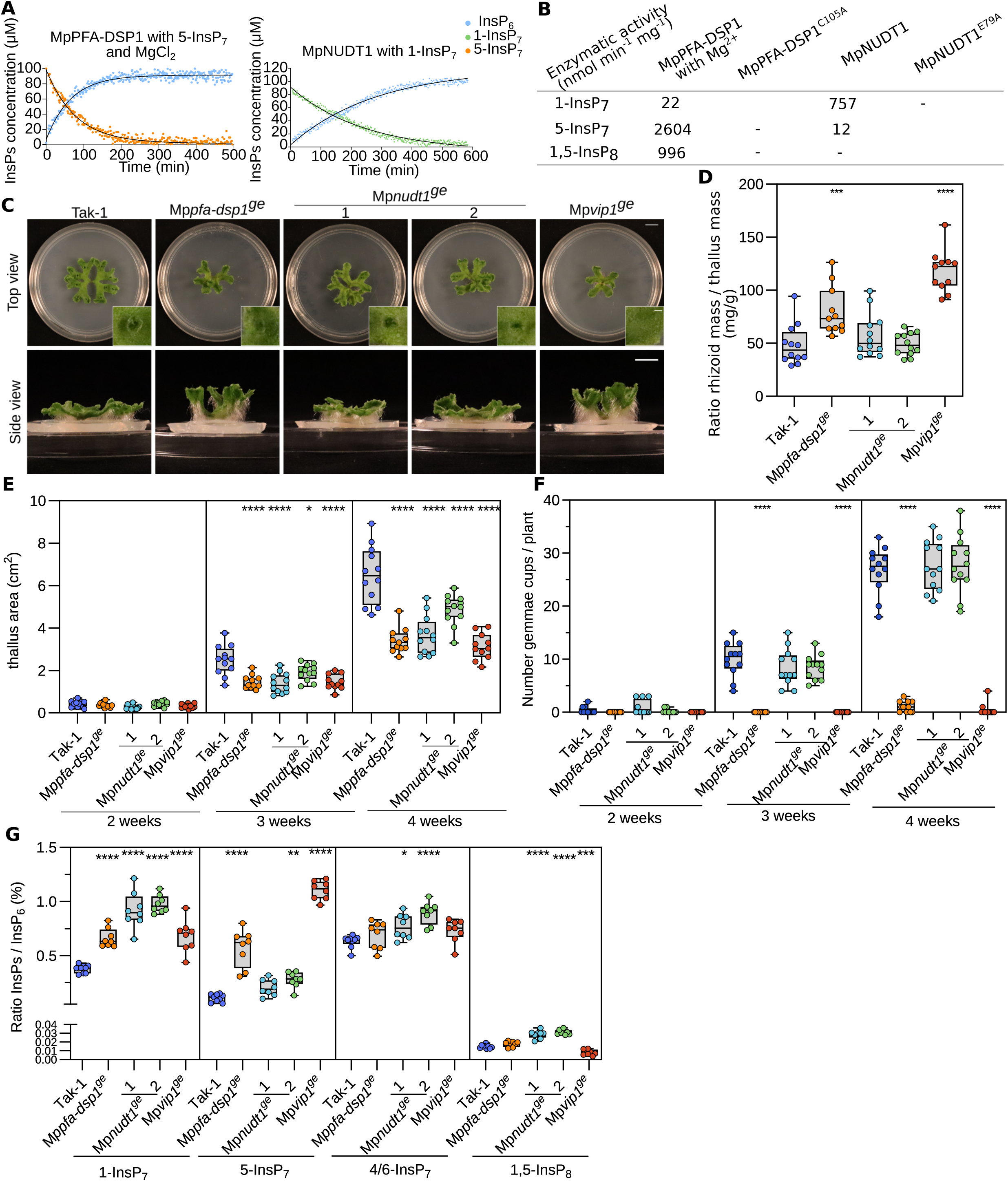
Inositol pyrophosphate phosphatases regulate Marchantia growth, development, and PP-InsP pools. **(A)** Pseudo-2D spin-echo difference NMR time course experiments for MpPFA-DSP1 and MpNUDT1 inositol phosphatase activities, using 100 μM of [^13^C_6_]5-InsP_7_ or [^13^C_6_]1-InsP_7_ as substrate, respectively. **(B)** Table summaries of the enzymatic activities of MpPFA-DSP1 and MpNUDT1 vs. PP-InsPs substrates. **(C)** Representative top and side views of 4-week-old Tak-1, Mp*pfa-dsp1^ge^*, Mp*nudt1^ge^* and Mp*vip1^ge^*mutant lines with different angles. Plants were grown from gemmae on ^½^B5 plates in continuous light at 22°C. Scale bar = 1 cm. Single gemmae cups are shown alongside, scale bar = 0.1 cm. **(D)** Thallus surface areas of Tak-1, Mp*pfa-dsp1^ge^*, Mp*nudt1^ge^* and Mp*vip1^ge^*mutant lines in time course experiments. Plants were grown from gemmae on ^½^B5 plates in continuous light with 22°C and one plant per round Petri dish as shown in **(C)**. For each genotype, 12 plants were taken. Statistical significance was assessed with a Dunnett test with Tak-1 as reference at each time point (**** p < 0.001, *** p < 0.005, ** p < 0.01, * p < 0.05). **(E)** Number of gemmae cups as a function of time for Tak-1, Mp*pfa-dsp1^ge^*, Mp*nudt1^ge^* and Mp*vip1^ge^*. Statistical significance was assessed with a Dunnett test with Tak-1 as reference at each time point (**** p < 0.001, *** p < 0.005, ** p < 0.01, * p < 0.05). **(F)** Rhizoids mass normalized to thallus mass of 4-week-old Tak-1, Mp*pfa-dsp1^ge^*, Mp*nudt1^ge^* and Mp*vip1^ge^*plants. Rhizoids were manually peeled with forceps. The weight of rhizoid was normalized by the thallus weight of the same plant. Statistical significance was assessed with a Dunnett test with Tak-1 as reference (**** p < 0.001, *** p < 0.005, ** p < 0.01, * p < 0.05). **(G)** PP-InsPs levels of 3-week-old Tak-1, Mp*pfa-dsp1^ge^*, Mp*nudt1^ge^* and Mp*vip1^ge^* plants. PP-InsPs were extracted with titanium oxide beads and then quantified by CE-ESI-MS. Data was normalized to the respective levels of InsP_6_.

### Deletion of Mp*PFA-DSP1*, Mp*NUDT1* or Mp*VIP1* alters cellular PP-InsP level, growth and development

Next, we generated Mp*pfa-dsp1^ge^* (nomenclature according to ref. (Bowman et al., 2016)) and Mp*nudt1^ge^* knockout mutants using CRISPR/Cas9 gene editing in *M. polymorpha* Tak-1 (Takaragaike-1) background (Supplementary Figure 7). For comparison, we also generated a Mp*vip1^ge^* (Mp8g06840) loss-of-function mutant, targeting the only PPIP5K gene in *M. polymorpha* (Supplementary Figure 7). 4-week-old Mp*pfa-dsp1^ge^* plants grown from gemmae exhibited a vertical thallus growth phenotype, a decreased thallus surface area, increased rhizoid mass and reduced number of gemma cups, when compared to Tak-1 (Figure 3C). Mp*vip1^ge^* mutants displayed similar phenotypes, while two independent CRISPR/Cas9 knockout alleles of Mp*nudt1^ge^* (Supplementary Figure 7) had only mild growth phenotypes (Figure 3C). In time course experiments, Mp*pfa-dsp1^ge^*, Mp*nudt1^ge^* and Mp*vip1^ge^*showed significantly reduced thallus surface areas (Figure 3D). Mp*pfa-dsp1^ge^* and Mp*vip1^ge^* but not Mp*nudt1^ge^* mutants had a strongly reduced number of gemma cups (Figure 3E). Mp*pfa-dsp1^ge^* and Mp*vip1^ge^* mutants showed increased rhizoid mass compared to Tak-1 (Figure 3F). Taken together, deletion of Mp*PFA-DSP1*, Mp*NUDT1* or Mp*VIP1* affects growth and development in *M. polymorpha*, with the Mp*pfa-dsp1^ge^*and Mp*vip1^ge^* mutants having rather similar phenotypes.

Quantification of PP-InsP levels in Tak-1 revealed that *M. polymorpha* contains levels of 1-InsP_7_, 5-InsP_7_, 1,5-InsP_8_ comparable to those found in Arabidopsis (Supplementary Figure 5), as well as the recently reported 4/6-InsP_7_ isomer (Riemer et al., 2021) (Figure 3G and Supplementary Figure 8A). Deletion of the PPIP5K MpVIP1 increases cellular 5-InsP_7_ pools, while decreasing 1,5-InsP_8_ levels, consistent with the enzymatic properties of the PPIP5K kinase domain (Wang et al., 2011) (Figure 3G). Mp*pfa-dsp1^ge^* mutants show increased 1-InsP_7_ and 5-InsP_7_ pools and wild-type-like 1,5-InsP_8_ levels (Figure 3G). Mp*nudt1^ge^*lines show an increase for 1-InsP_7_ consistent with the preferred *in vitro* substrate of MpNUDT1 (Figure 3A, B, G). 1,5-InsP_8_ levels are higher in Mp*nudt1^ge^* when compared to Tak-1 (Figure 3G). None of the mutants affected the levels of 4/6- InsP_7_, suggesting that its biosynthesis/catabolism may not be catalyzed by MpVIP1, MpPFA-DSP1 or MpNUDT1 in *M. polymorpha* (Figure 3G). Together, Marchantia VIP1, PFA-DSP1 and NUDT1 are *bona fide* PP-InsP metabolizing enzymes *in vitro* and *in planta*.

### MpPFA-DSP1, MpNUDT1 and MpVIP1 regulate Pi homeostasis in Marchantia

Arabidopsis *vih1 vih2* mutants with reduced 1,5-InsP_8_ pools overaccumulate cellular Pi (Ried et al., 2021; Zhu et al., 2019). Consistently, Mp*pfa-dsp1^ge^* and Mp*nudt1^ge^* plants with increased 1,5-InsP_8_ levels (Figure 3G) have associated lower Pi concentrations (Figure 4A). Our Mp*vip1^ge^* mutant has lower 1,5-InsP_8_ concentrations (Figure 3G) but Pi levels not significantly different from the Tak-1 control (Figure 4A). RNA-seq analysis comparing Tak-1 plants grown under Pi-sufficient vs. Pi-starved conditions revealed that only Mp*NUDT1* expression is repressed under Pi starvation (Figure 4B). We could not detect _pro_Mp*PFA-DSP1* or _pro_Mp*NUDT1* promoter activity in promoter::GUS fusions, whereas _pro_MpVIP1 showed a robust signal under both Pi-sufficient and Pi starvation conditions (Supplementary Figure 9). We compared Tak-1 plants grown under Pi-sufficient and Pi-starved conditions by RNA-seq to define PSI marker genes (Figure 4C), some of which are orthologs of the known PSI genes in Arabidopsis and in other plant species (Cuyas et al., 2023). Next, we analyzed PSI marker gene transcript levels in our Mp*pfa-dsp1^ge^*, Mp*nudt1^ge^*and Mp*vip1^ge^* mutants with Tak-1 grown under Pi sufficient conditions. We found that Mp*SPX* (Mp1g27550) transcript levels were decreased in Mp*pfa-dsp1^ge^* and in Mp*vip1^ge^*. Mp*PHO1;H4* (Mp4g19710) levels were decreased in Mp*pfa-dsp1^ge^*and Mp*vip1^ge^* and increased in Mp*nudt1^ge^* (Figure 4D). Pi transporter Mp*PHT1;4* (Mp2g20620) transcript levels are higher in our Mp*pfa-dsp1^ge^*and Mp*vip1^ge^* mutants when compared to Tak-1 (Figure 4D). Taken together, these experiments support a function for PP-InsPs in *M. polymorpha* Pi homeostasis, with the Mp*pfa-dsp1^ge^* and Mp*vip1^ge^*mutants showing similar gene expression patterns (Figure 4E). However, manually curated gene ontology analyses of the differentially expressed genes (DEGs) revealed that PSI genes only represent a small pool of the total DEGs (Figure 4E).

**Figure 4.**
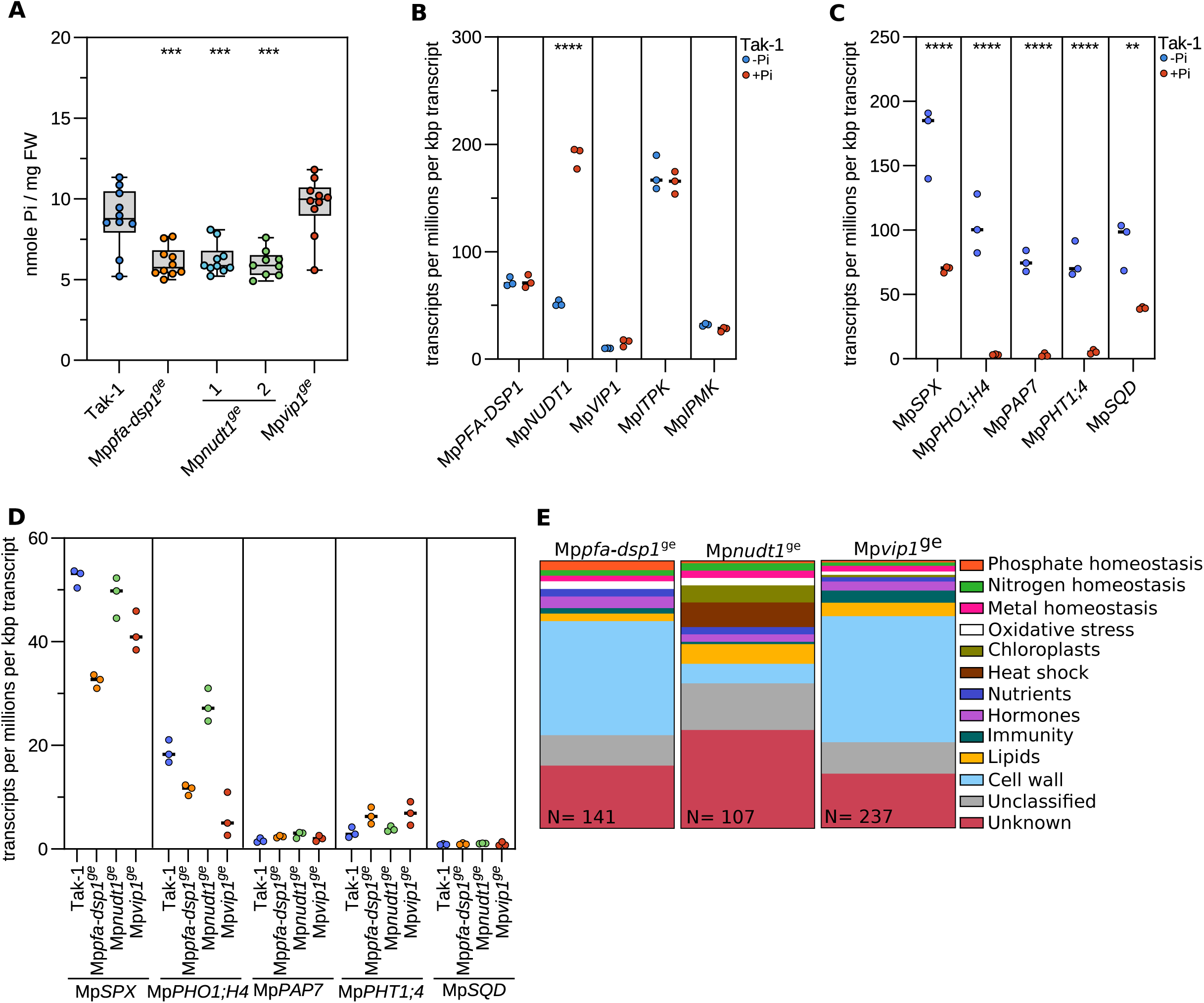
PSI gene expression and Pi homeostasis are affected in Mp*pfa-dsp1^ge^ and* Mp*nudt1^ge^*mutants. **(A)** Total Pi levels of 3-week-old Tak-1, Mp*pfa-dsp1^ge^*, Mp*nudt1^ge^* and Mp*vip1^ge^* plants grown under Pi-sufficient conditions. Technical triplicates were done for the standards and duplicates for all samples. Statistical significance was assessed with a Dunnett test with Tak-1 as reference (**** p < 0.001, *** p < 0.005, ** p < 0.01, * p < 0.05). **(B)** Quantification of the PP-InsP-metabolizing MpPFA-DSP1, MpNUDT1, MpVIP1, MpITPK1 and MpIPMK enzyme transcripts from RNA-seq experiments performed on 2-week-old Tak-1 plants grown in either no phosphate (-Pi) or in 0.5 mM K_2_HPO_4_/KH_2_PO_4_ (+Pi). Counts were normalized by the number of reads in each dataset and by the length of each transcript. **(C)** Identification of PSI marker in *Marchantia polymorpha* comparing 2-week-old Tak-1 plants grown in -Pi and +Pi conditions as in **(B)**. **(D)** Gene expression of the PSI marker genes defined in **(C)** comparing 3-week-old Tak-1, Mp*pfa-dsp1^ge^*, Mp*nudt1^ge^* and Mp*vip1^ge^* grown under Pi-sufficient conditions to Tak-1. **(E)** Manually curated gene-ontology classification of DEGs of 3-week-old Mp*pfa-dsp1^ge^*, Mp*nudt1^ge^*and Mp*vip1^ge^* mutant lines vs. Tak-1. DEGs with | log_2_(FC)|>2 and p < 0.05 were considered differentially expressed.

### Cell wall composition is altered in Mp*pfa-dsp1^ge^* and Mp*vip1^ge^* mutants

The large number of DEGs unrelated to Pi homeostasis prompted us to investigate other pathways potentially affected by the altered PP-InsP levels in Mp*pfa-dsp1^ge^*, Mp*nudt1^ge^* and Mp*vip1^ge^*. We selected metal ion and cell wall homeostasis for further analysis (Figure 4E). Several metal ion transporters, metallothioneins and oxidoreductases are differentially expressed in our PP-InsP enzyme mutants (Supplementary Figure 10A), but we did not observe unique, ion-specific differences in the ionomic profiles of Mp*pfa-dsp1^ge^*, Mp*nudt1^ge^* and Mp*vip1^ge^* mutants compared to Tak-1 (Supplementary Figure 10B, C). Rather, Mp*pfa-dsp1^ge^* and Mp*vip1^ge^* appear to contain slightly elevated concentrations of various mono- and divalent cations, including potassium, magnesium, calcium, zinc and molybdenum (Supplementary Figure 10B, C).

The largest set of DEGs in our Marchantia RNA-seq experiments maps to cell wall related genes, particularly to a large number of class III peroxidases (Figure 5A) (Almagro et al., 2009). Notably, *AtPFA-DSP2* OX lines also show altered gene expression patterns for many cell wall related genes, including peroxidases (Figure 5B). High peroxidase activity has been previously reported from *M. polymorpha* cell wall fractions (Ishida et al., 1985), and therefore we investigated cell wall related phenotypes in our different mutants. Ruthenium red-stained transverse cross sections of 3-week-old thalli revealed increased staining in the dorsal and ventral epidermides of Mp*pfa-dsp1^ge^* and Mp*vip1^ge^*mutants, when compared to Tak-1 (Figure 5C-E), indicating increased acidic pectin levels in these two mutants. Fluorol yellow staining for lipidic compounds such as suberin or cutin also showed strong signals in the dorsal and ventral epidermal layers (Figure 5E), suggesting that the Mp*pfa-dsp1^ge^* and Mp*vip1^ge^*mutants may contain higher levels of polyester cell wall polymers. In contrast, Renaissance SR2200 (which stains cellulose, hemicellulose and callose) revealed a uniform staining pattern across all mutants analyzed (Figure 5E), suggesting that only specific cell wall components are altered in our Mp*pfa-dsp1^ge^* and Mp*vip1^ge^*lines. Taken together, our transcriptomic and histological analyses indicate cell wall composition changes in the Mp*pfa-dsp1^ge^* and Mp*vip1^ge^* epidermal layers.

**Figure 5.**
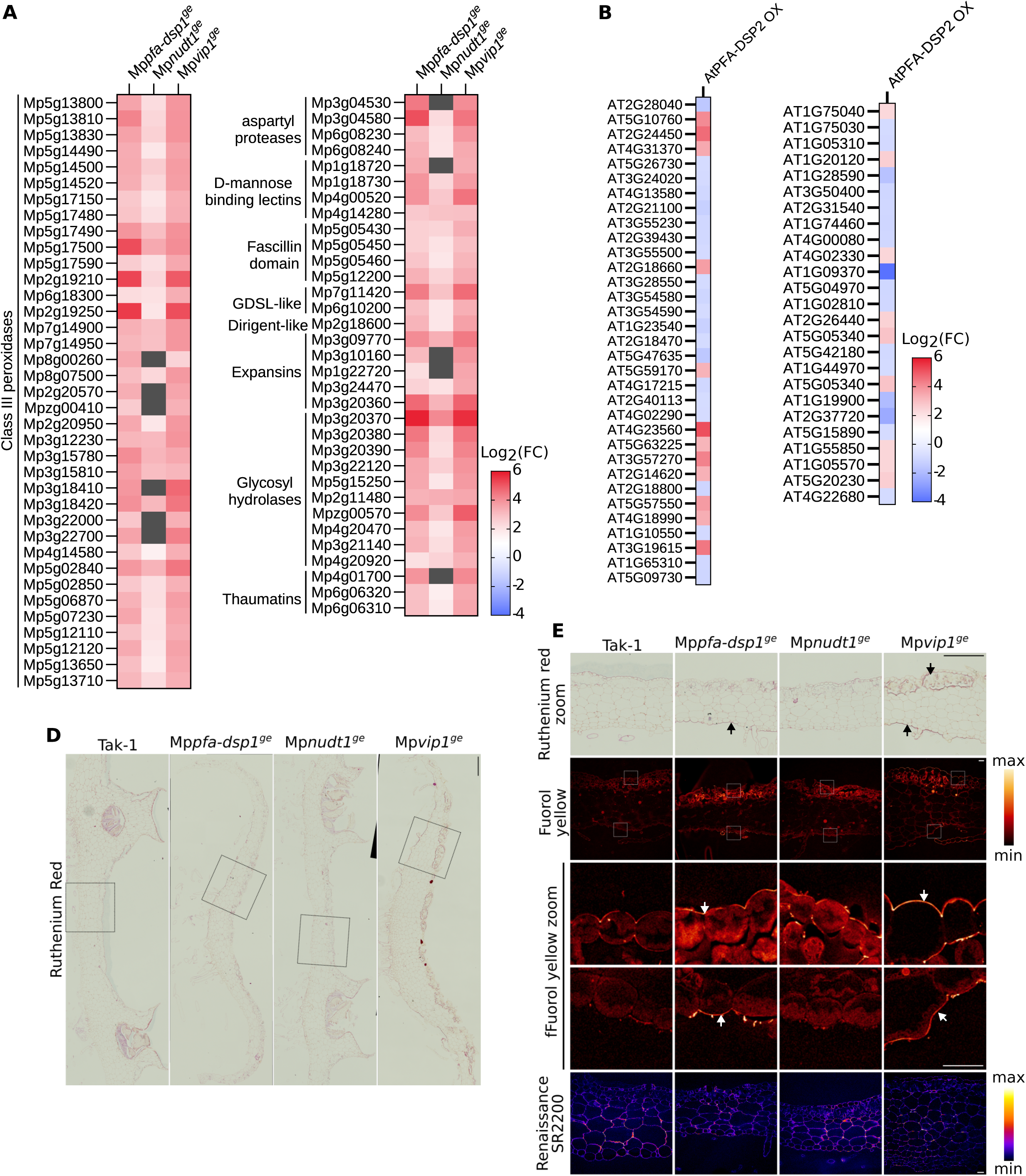
Cell wall composition is altered in Mp*pfa-dsp1^ge^* and Mp*vip1^ge^* mutant plants. **(A)** Heatmap of DEGs in 3-week-old Mp*pfa-dsp1^ge^*, Mp*nudt1^ge^*and Mp*vip1^ge^* plants grown under Pi-sufficient conditions vs. Tak-1. Known marker genes significantly different from Tak-1 and putatively involved in cell wall homeostasis are displayed. Grey boxes = not differentially expressed. **(B)** Heatmap of DEGs of 2-week-old At*PFA-DSP2* OX plants vs. Col-0. **(C)** Schematic representation of a transversal thallus cross section of *Marchantia polymorpha*. **(D)** Fixed transverse cross-sections at the level of gemmae cups from 3-week-old Tak-1, Mp*pfa-dsp1^ge^*, Mp*nudt1^ge^* and Mp*vip1^ge^* plants, stained with ruthenium red. **(E)** From top to bottom: Enlarged view of the ruthenium red-stained sections from **(D)** (scale bar=500 μm), fluorol yellow, enlarged view of fluorol yellow-stained dorsal side, enlarged view of fluorol yellow-stained ventral side (scale bar=10 μm), total view of the Renaissance SR2200-stained cross section (scale bar=50 μm). Look-up tables for fluorol yellow and Renaissance SR2200 are shown alongside, regions in Mp*pfa-dsp1^ge^*or Mp*vip1^ge^* enriched in cell wall material compared to Tak-1 are marked by arrows.

### Deletion of the VIP1 phosphatase domain in Mp*fa-dsp1^ge^* affects plant growth and nitrogen accumulation

Based on the similar growth phenotypes, PP-InsP pools, gene expression changes and cell wall defects of our Mp*pfa-dsp1^ge^* and Mp*vip1^ge^* mutants, we next performed genetic interaction studies between Mp*PFA-DSP1* and Mp*VIP1*. Since MpVIP1 is a bifunctional enzyme with both PP-InsP kinase and phosphatase activity, we targeted the C-terminal histidine acid phosphatase domain (PD) in MpVIP1 by CRISPR/Cas9-mediated gene editing. The resulting Mp*vip1Δpd^ge^* mutant lacks the C-terminal phosphatase domain while retaining the N-terminal PPIP5K kinase domain (Figure 6A and Supplementary Figure 7). We also isolated a Mp*pfa-dsp1^ge^* Mp*vip1Δpd^ge^*double mutant (Figure 6A). Notably, we could not recover Mp*nudt1^ge^* Mp*pfa-dsp1^ge^*or Mp*nudt1^ge^* Mp*vip1Δpd^ge^* double mutants, potentially indicating that these mutant combinations are not viable, as in yeast (Sanchez et al., 2023, 2019). 4-week-old Mp*vip1Δpd^ge^* plants grown from gemmae had reduced thallus surface areas when compared to Tak-1 (Figure 6B). Interestingly, Mp*vip1Δpd^ge^* and Mp*vip1^ge^* mutants display similar growth phenotypes (Figure 6A, B). Thallus size is reduced further in the Mp*pfa-dsp1^ge^* Mp*vip1Δpd^ge^*double mutants, suggesting that inositol 1- and 5-pyrophosphate phosphatase activities are required for *M. polymorpha* growth and development (Figure 6A, B). Notably, the Mp*pfa-dsp1^ge^*, Mp*nudt1^ge^* and Mp*pfa-dsp1^ge^*Mp*vip1Δpd^ge^* mutants accumulate less Pi compared to Tak-1, when grown under Pi sufficient conditions (Figure 6C, compare Figure 4a).

**Figure 6.**
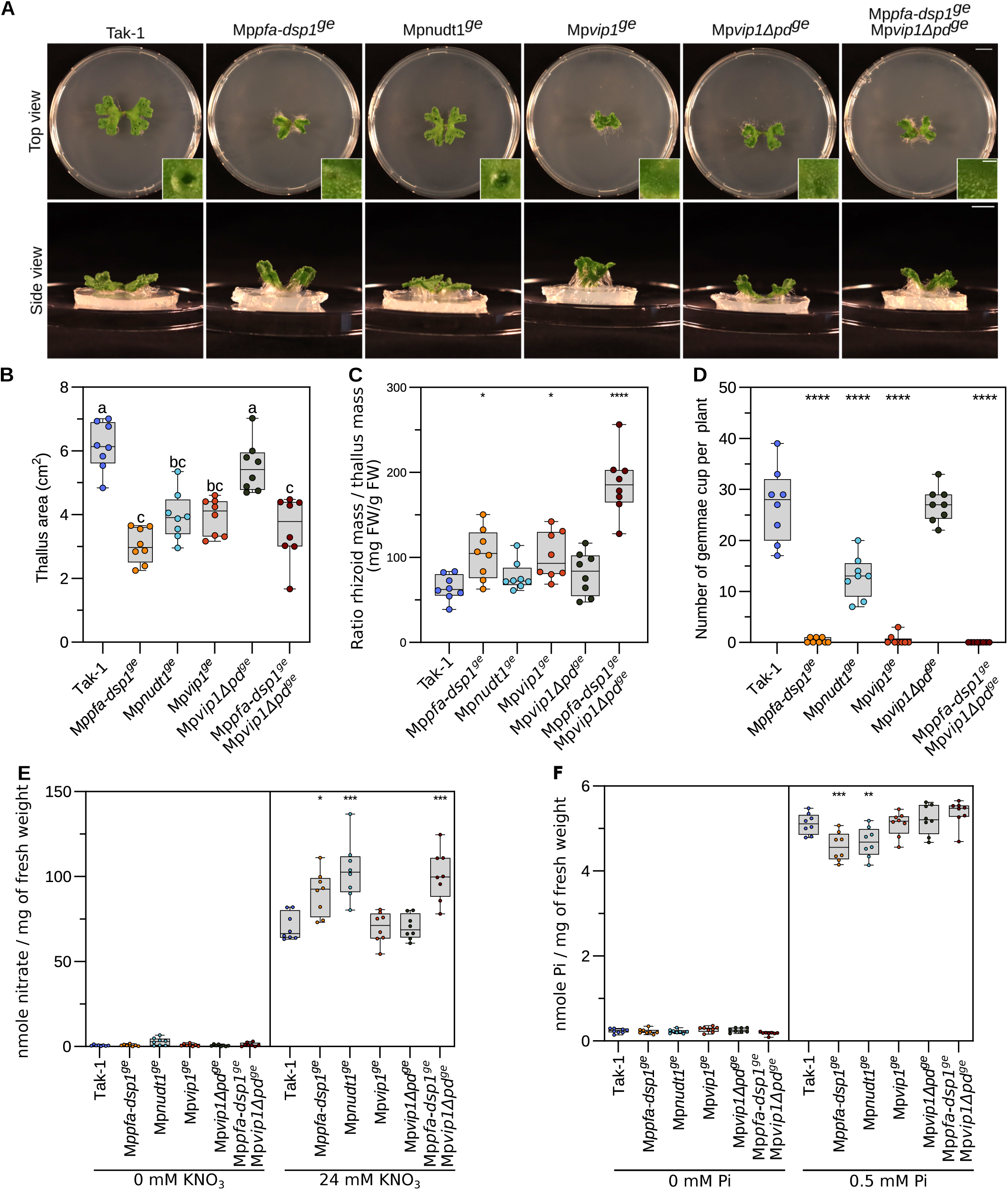
PP-InsP catabolic enzymes regulate Pi and nitrate homeostasis in Marchantia. **(A)** Growth phenotypes of 3-week-old Tak-1, Mp*pfa-dsp1^ge^*, Mp*nudt1^ge^* and Mp*vip1^ge^*, Mp*vipΔpd^ge^*and Mp*vip1^ge^* Mp*vipΔpd^ge^* plants. Plants were grown from gemmae on ^½^B5 plates in continuous light at 22°C. Scale bar = 1 cm. Single gemmae cups are shown alongside, scale bar = 0.1 cm. **(B)** Quantification of projected thallus surface areas of 3-week-old Tak-1, Mp*pfa-dsp1^ge^*, Mp*nudt1^ge^*, Mp*vip1^ge^*, Mp*vipΔpd^ge^* and Mp*vip1^ge^* Mp*vipΔpd^ge^* plants. Tukey-type all-pairs comparisons between the genotypes (Tukey et al., 1985) were performed in the R package multcomp (Hothorn et al., 2008). **(C)** Number of gemmae cups of 4-week-old Tak-1, Mp*pfa-dsp1^ge^*, Mp*nudt1^ge^* and Mp*vip1^ge^* plants. Statistical significance was assessed with a Dunnett test with Tak-1 as reference (**** p < 0.001, *** p < 0.005, ** p < 0.01, * p < 0.05). **(D)** Rhizoids mass normalized to thallus mass of 4-week-old Tak-1, Mp*pfa-dsp1^ge^*, Mp*nudt1^ge^*, Mp*vip1^ge^*, Mp*vipΔpd^ge^* and Mp*vip1^ge^* Mp*vipΔpd^ge^* plants. Rhizoids were manually peeled with forceps. The weight of rhizoid was normalized by the thallus weight of the same plant. Statistical significance was assessed with a Dunnett test with Tak-1 as reference (**** p < 0.001, *** p < 0.005, ** p < 0.01, * p < 0.05). **(E)** Nitrate quantification of 2-week-old Tak-1, Mp*pfa-dsp1^ge^*, Mp*nudt1^ge^*, Mp*vip1^ge^*, Mp*vipΔpd^ge^*and Mp*vip1^ge^* Mp*vipΔpd^ge^* plant lines grown under nitrate starvation or control conditions. 8 plants were used per genotype. Nitrate was quantified adapting the Miranda, spectrophotometric method (Miranda et al., 2001). Technical triplicates were done for the standards and duplicates for all samples. Statistical significance was assessed with a Dunnett test with Tak-1 as reference (**** p < 0.001, *** p < 0.005, ** p < 0.01, * p < 0.05). **(F)** Total Pi levels of 2-week-old Tak-1, Mp*pfa-dsp1^ge^*, Mp*nudt1^ge^*, Mp*vip1^ge^*, Mp*vipΔpd^ge^* and Mp*vip1^ge^*Mp*vipΔpd^ge^* plants grown under Pi-starvation or Pi-sufficient (0.5 mM K_2_HPO_4_/KH_2_PO_4_) conditions. Technical triplicates were done for the standards and duplicates for all samples. Statistical significance was assessed with a Dunnett test with Tak-1 as reference (**** p < 0.001, *** p < 0.005, ** p < 0.01, * p < 0.05). **(D)**

It has been previously reported that SPX domains are regulators of nitrate signaling in rice and in Arabidopsis (Hu et al., 2019; Ueda et al., 2020; Zhang et al., 2021). Therefore, we quantified nitrate levels in 4-week-old plants grown on regular B5 medium (see Methods). Under these nitrate-sufficient growth conditions, all of our mutants accumulate nitrate to higher levels compared to Tak-1, with the Mp*pfa-dsp1^ge^* Mp*vip1Δpd^ge^* double mutant having the strongest effect (Figure 6D). This suggests, that PP-InsPs may affect nitrate homeostasis in *M. polymorpha*, although it is unclear which PP-InsP isomer may be involved (Figure 3G and 6D). Taken together, the Mp*pfa-dsp1^ge^* Mp*vip1Δpd^ge^* double mutant phenotypes suggests that inositol 1- and 5-pyrophosphate phosphatase activities regulate *M. polymorpha* growth and development.

## Discussion

Important physiological functions for inositol pyrophosphates in Arabidopsis have been highlighted by analysis of ITPK and PPIP5K loss-of-function mutants (Dong et al., 2019; Laha et al., 2019, 2015; Riemer et al., 2021; Zhu et al., 2019). Genetic, quantitative biochemical and structural evidence support a role for the 1,5-InsP_8_ isomer as an essential nutrient messenger in Arabidopsis Pi homeostasis (Dong et al., 2019; Guan et al., 2022; Ried et al., 2021; Wild et al., 2016; Zhu et al., 2019), similar to that described in yeast (Chabert et al., 2023) and human (Li et al., 2020). Under Pi sufficient growth conditions, high cellular ATP/ADP ratios support 1,5-InsP _8_ biosynthesis by activating the PPIP5K kinase domain (Gu et al., 2017; Zhu et al., 2019). At the same time, cellular Pi acts as an inhibitor of the PPIP5K histidine acid phosphatase domain, resulting in a net accumulation of the 1,5-InsP_8_ nutrient messenger (Gu et al., 2017; Zhu et al., 2019). Under Pi starvation conditions, ATP and Pi levels decrease, inhibiting the kinase and stimulating the inositol 1-pyrophosphatase activity of PPIP5Ks (Gu et al., 2017; Zhu et al., 2019). However, plants harboring phosphatase-dead versions of the PPIP5K AtVIH2 did not show Pi homeostasis-related phenotypes (Zhu et al., 2019), suggesting that other PP-InsP phosphatases may be involved in 1,5-InsP_8_ catabolism in Arabidopsis.

Here, we characterize three PFA-DSP-type and three NUDIX-type enzymes as inositol pyrophosphate phosphatases in Arabidopsis. Previous studies (Gaugler et al., 2022; Wang et al., 2022) and our biochemical analysis reveal AtPFA-DSPs as specific inositol 5-pyrophosphate phosphatases. As in the case of yeast Siw14 (Steidle et al., 2016), 5-InsP_7_ is the preferred *in vitro* substrate for AtPFA-DSP1 in the presence and absence of Mg^2+^ ions (Wang et al., 2022; Kurz et al., 2023) (Figure 1A, B and Supplementary Figure 2). The MpPFA-DSP1 ortholog shares the substrate specificity and overall activity with the Arabidopsis enzyme (Figure 3B and Supplementary Figure 6). We found that 5-InsP_7_ is the preferred *in vitro* substrate for AtNUDT17 (Figure 1A, B and Supplementary Figure 2). However, the enzyme is much less active compared to AtPFA-DSP1 (Figure 1A, B). In contrast to AtNUDT17, MpNUDT1 strongly prefers 1-InsP_7_ as substrate *in vitro* and *in vivo* (Figure 3A, B, G). A preference for different pyrophosphorylated substrates has been previously described for yeast and human NUDIX enzymes (Márquez-Moñino et al., 2021; Zong et al., 2021). The fact that AtNUDT13 can also hydrolyze 1- and 5-InsP_7_ (Supplementary Figure 2B, C) suggests that we could recover some but not all PP-InsP phosphatases in our 5PCP-InsP _5_ interaction screen (Supplementary Figure 1A, B) (Furkert et al., 2020; Wu et al., 2016).

Overexpression of *AtPFA-DSP1*, *AtPFA-DSP2* or *AtPFA-DSP4* resulted in stunted growth phenotypes associated with a reduction in 5-InsP_7_ and 1,5-InsP_8_ levels (Figure 1C-F and Supplementary Figure 4). Pi levels are elevated in *AtPFA-DSP2* OX lines and PSI gene expression is strongly upregulated (Figure 2C, E). Since AtPFA-DSP1 and AtNUDT17 have a similar substrate preference *in vitro* and *in vivo* (Figure 1B, F), we speculate that the weaker overexpression effects in our *AtNUDT17* OX lines (Figure 1C and Supplementary Figure 4) are related to the lower enzyme activity of AtNUDT17 (Figure 1B). Consistent with our study, overexpression of *AtPFA-DSP1* in tobacco and in Arabidopsis resulted in reduced InsP_7_ pools (Gaugler et al., 2022).

Our *nudt17/18/21* loss-of-function mutants are indistinguishable from wild type and show only minor changes in PP-InsP accumulation and repression of PSI gene expression (Figure 1C, F, 2D, E). Since all three NUDT enzymes are expressed at seedling stage (Figure 2A), we speculate that other NUDT family members such as AtNUDT13 (Supplementary Figure 2B, C) may act redundantly with AtNUDT17/18/21 in PP-InsP catabolism. Although no loss-of-function phenotypes for NUDT enzymes were observed, their induction under Pi starvation conditions suggests that these PP-InsP phosphatases may regulate Pi homeostasis in Arabidopsis (Figure 2A, B).

*M. polymorpha* contains 9 PFA-DSP and 20 NUDT genes. We were able to define loss-of-function phenotypes for Mp*pfa-dsp1^ge^*and Mp*nudt1^ge^* single mutants. Overall, both Mp*pfa-dsp1*^ge^ and Mp*nudt1*^ge^ show reduced thallus growth rates in time course experiments (Figure 3C, D). To our surprise, Mp*vip1^ge^* plants (originally generated as a control) shared the vertical thallus growth phenotype, a smaller thallus surface area, increased rhizoid mass, and reduced number of gemma cups with Mp*pfa-dsp1^ge^* (Figure 3C-F). Similar phenotypes have been reported previously for PIN auxin transporter overexpression lines, and for auxin response factor loss-of-function mutants in *M. polymorpha (Kato et al., 2017; Tang et al., 2024)*. Notably, loss-of-function mutants of the 5-InsP_7_ synthesizing ITPK1 kinase show altered auxin responses in Arabidopsis (Laha et al., 2022). A PP-InsP binding site has been previously identified in the auxin receptor AtTIR1 (Sheard et al., 2010), which may sense the AtITPK1 reaction product (Laha et al., 2022). Thus, auxin responses could be altered in Mp*pfa-dsp1^ge^* and Mp*vip1^ge^* plants, which contain higher 5-InsP_7_ levels (Figure 3G). However, our RNA-seq analyses did not reveal any major changes in the expression of auxin-regulated genes (Figure 4E). Changes in PP-InsP levels alter Pi homeostasis in Arabidopsis (Gaugler et al., 2022; Riemer et al., 2021; Zhu et al., 2019), and therefore we characterized PP-InsP concentrations and Pi starvation-related phenotypes in our different phosphatase loss-of-function mutants. 1- and 5-InsP_7_ levels are increased in Mp*pfa-dsp1^ge^* mutants compared to Tak-1, while 1,5-InsP_8_ concentrations are only slightly increased (Figure 3G). Both Mp*nudt1^ge^* alleles overaccumulate 1-InsP_7_ and 1,5-InsP_8_ (Figure 3G). Taken together, PFA-DSP and NUDT enzymes in Marchantia and in Arabidopsis contribute to PP-InsP catabolism.

We observed that in contrast to the Arabidopsis *vih1 vih2* mutant (Zhu et al., 2019), Mp*vip1^ge^* plants are viable (Figure 3C) and do not overaccumulate phosphate under Pi-sufficient growth conditions (Figure 4A, 6C). To our knowledge, Mp*VIP1* is a single-copy gene in *M. polymorpha.* The Mp*vip1^ge^* mutant contains lower levels of 1,5-InsP_8_ when compared to Tak-1 (Figure 3G). However, for several PSI marker genes identified in our RNA-seq experiments (Figure 4C, see also ref. (Rico-Reséndiz et al., 2020)), we observed gene repression rather than constitutive activation in Mp*vip1^ge^* plants (Figure 4D). Consistent with this, Mp*vip1^ge^*phenocopies the Mp*pfa-dsp1^ge^* mutant, which also has higher 5-InsP_7_ levels but only slightly increased 1,5-InsP_8_ pools (Figure 3G), associated with PSI marker gene repression (Figure 4D). Deletion of the C-terminal histidine acid phosphatase in MpVIP1 (Mp*vip1Δpd^ge^*) resulted in reduced thallus growth, similar to the Mp*vip1^ge^* and Mp*pfa-dsp1^ge^*mutants (Figure 6A, B). This suggests that both the PPIP5K kinase and the histidine acid phosphatase activities contribute to this phenotype. In line with this, thallus size is further reduced in Mp*pfa-dsp1^ge^* Mp*vip1Δpd^ge^* double mutants (Figure 6A, B), suggesting that inositol 1- and 5-pyrophosphate phosphatase activities are required for normal growth and development in *M. polymorpha*. However, the Mp*vip1Δpd^ge^*mutant has wild-type-like Pi levels and the Mp*pfa-dsp1^ge^*Mp*vip1Δpd^ge^* double mutant had Pi levels similar to Mp*pfa-dsp1^ge^*(Figure 6C). Therefore, our data do not support an isolated function for MpVIP1 as master regulator of Marchantia Pi homeostasis, unlike what has been reported in Arabidopsis (Dong et al., 2019; Zhu et al., 2019). We speculate that *M. polymorpha* may contain a second, sequence-divergent PP-InsP kinase able to synthesize 1,5-InsP_8_. In line with this, vip1Δ (the single PPIP5K in baker’s yeast) mutants still contain detectable levels of 1,5-InsP_8_ (Chabert et al., 2023). Linking Mp*pfa-dsp1^ge^*, Mp*nudt1^ge^* or Mp*vip1^ge^*mutant phenotypes to isomer-specific PP-InsP level changes is complicated by compensatory changes in gene expression for other PP-InsP metabolizing enzymes, as indicated by our RNA-seq experiments (Supplementary Figure 8B). Importantly, other PFA-DSP and NUDT-type inositol pyrophosphate phosphatases may exist in *M. polymorpha*.

Based on our RNA-seq analyses (Figure 4E), we additionally quantified cell wall-related phenotypes in the different mutant backgrounds (Figure 5D, E). Indeed, gene expression changes for many cell wall-related and carbohydrate-active enzymes could be associated with specific changes in cell wall composition in the Mp*pfa-dsp1^ge^* and Mp*vip1^ge^* mutants (Figure 5D, E). It has been previously reported that Pi starvation induces cellulose synthesis (Khan et al., 2024), and that ectopic overexpression of wheat VIH2 in Arabidopsis resulted in higher cellulose, arabinoxylan and arabinogalactan levels (Shukla et al., 2021). Similarly, extracellular Pi sensing has been associated with callose deposition in the root tip (Balzergue et al., 2017; Müller et al., 2015). Our work suggests that altered PP-InsP levels in the Mp*pfa-dsp1^ge^* and Mp*vip1^ge^* mutants induce changes in Marchantia cell wall composition.

All our mutants show increased nitrate levels (Figure 6D), with the Mp*pfa-dsp1^ge^* Mp*vip1Δpd^ge^* double mutant having the strongest effect. Consistent with this, nitrogen homeostasis-related genes are differentially expressed in Mp*pfa-dsp1^ga^*, Mp*nudt1^ge^* and Mp*vip1^ge^* (Figure 4E), and nitrate transporters are induced in our *AtPFA-DSP2* OX and in *AtNUDT17* OX lines (Figure 2E). SPX inositol pyrophosphate receptors (Wild et al., 2016) have previously been implicated in nitrogen sensing and signaling (Hu et al., 2019; Zhang et al., 2021; Ueda et al., 2020; Kant et al., 2011; Liu et al., 2017). A genetic interaction between VIP1 and nitrogen starvation has been reported in *Chlamydomonas reinhardtii* (Couso et al., 2016). Our PP-InsP catabolic mutants now suggest a more direct link between cellular PP-InsP pools and nitrate homeostasis (Figure 3G, 6D). Interestingly, alterations in nitrogen supply affect cell wall organization and composition in several plant species (Fernandes et al., 2013; Rivai et al., 2021; Głazowska et al., 2019), providing an alternative rationale for the cell wall defects observed in our Mp*pfa-dsp1^ge^*and Mp*vip1^ge^* mutants (Figure 5). Future studies will elucidate the molecular mechanisms linking PP-InsPs with plant nitrogen homeostasis, and with cell wall architecture.

In conclusion, all three families of inositol pyrophosphate phosphatases present in plants contribute to the control of cellular PP-InsP pools, and changes in these pools regulate growth as well as phosphate, nitrogen and cell wall homeostasis in Marchantia.

## Methods

### Plant material

*Arabidopsis thaliana* ecotype Col-0, *nudt17/18/21*, AtPFA-DSP1, 2 or 4 OX, and AtNUDT17, 18 or 21 OX lines, and the previously reported *phr1 phl1* (Bustos et al., 2010), *vih1 vih2 phr1 phl1* (Zhu et al., 2019), and *pho2-1* (Delhaize and Randall, 1995) lines were gas sterilized, and after 2 d of stratification on ^½^MS (1.4 g/L MS basal salt mixture, 0.1 g/L MES, pH 5.7, plant agar= 8 g/L) grown for one week at 22°C and in 18 h / 6h light / dark cycles. Seedlings were transferred to soil and for rosette size quantification images were taken from 3-week-old plants. Wild type and CRISPR/Cas9-gene edited *Marchantia polymorpha* plants were Takaragaike-1 (Tak-1) males (Ishizaki et al., 2008). Plants were asexually maintained and propagated through gemmae growth on ½ Gamborg B5 medium (Sigma) adjusted to pH 5.5 with KOH, under constant LED-source white light (60 μmol/m^2^/s) at 22°C on 90 mm square Petri dishes (Greiner) containing 0.8 % (w/v) plant cell culture agar (Huber lab).

### 5PCP-InsP5 pull-down assay

Pull-down assays were performed with either resin-immobilized 5PCP-InsP_5_ or Pi, as previously described (Wu et al., 2016). *Arabidopsis thaliana* ecotype Col-0 seeds were germinated in ^1/2^MS agar plates for 5 d and transferred to liquid ^½^MS medium (containing 1 % [w/v] sucrose) in the presence of 0.2 µM (-Pi) or 1 mM (+Pi) K_2_HPO_4_/KH_2_PO_4_ (pH 5.7) for 10 d (Supplementary Figure 1A). Seedlings were collected, pat dry, frozen and ground to a fine powder in liquid N_2_. For each sample, 6-10 g of fresh tissue were incubated for 1 h on ice with a 1:3 ratio of extraction buffer (50 mM Tris-HCl pH 7.5, 150 mM NaCl, 1 mM EDTA, 10 % [v/v] glycerol, 5 mM dithiothreitol [DTT], 0.5 % [v/v] IGEPAL CA-630, 1 tablet of plant protease inhibitor cocktail [Roche] and 1 mM PMSF), with gentle shaking. Samples were then centrifuged at 16,000 x g for 20 min at 4°C, the supernatants were then filtered using Miracloth (Merck) and transferred to new Eppendorf tubes. Protein concentrations were measured using the Bradford assay and samples were diluted to a final concentration of 5 mg/mL. For each sample, 150 μL of beads slurry (Wu et al., 2016) was added to a new tube. Beads were pulled down by brief centrifugation at 100 x g and at 4°C and then washed twice with cold extraction buffer. The washed beads were then added to the protein extracts and incubated for 3 h in the cold room with gentle shaking. Eppendorf tubes were centrifuged at 100 x g for 30 s at 4°C, washed three times with extraction buffer, and eluted in 30 μL of elution buffer (50 mM Tris-HCl pH 7.5, 150 mM NaCl, 1 mM EDTA, 10 % [v/v] glycerol, 5 mM DTT and 20 mM InsP_6_) for 30 min in cold room with gentle shaking. Tubes were centrifuged and the supernatant was collected. A second elution was performed with an incubation of 30 μL of elution buffer overnight in the cold room with gentle shaking. The supernatant of this elution was collected and the two elutions were pooled. The remaining beads present in the eluate were removed by passing it through a Micro Bio-Spin chromatography column (Bio-Rad). 20 μL of 4x Laemmli sample buffer (Bio-Rad) was added, and samples were incubated at 95°C for 5 min. 10 μL of each sample was analyzed by SDS-PAGE followed by silver staining, the remaining sample was loaded on a 12 % mini polyacrylamide gel, migrated about 2 cm and stained by Coomassie. Gel lanes between 15-300 kDa were excised into 5-6 pieces and digested with sequencing-grade trypsin (Shevchenko et al., 2006). Extracted tryptic peptides were dried and resuspended in 0.05 % trifluoroacetic acid, 2 % (v/v) acetonitrile.

Data-dependent LC-MS/MS analyses of samples were carried out on a Fusion Tribrid Orbitrap mass spectrometer (Thermo Fisher Scientific) interfaced through a nano-electrospray ion source to an Ultimate 3000 RSLCnano HPLC system (Dionex). Peptides were separated on a reversed-phase custom packed 45 cm C18 column (75 μm ID, 100Å, Reprosil Pur 1.9 um particles, Dr. Maisch, Germany) with a 4-76% acetonitrile gradient in 0.1 % formic acid (total time 65 min). Full MS survey scans were performed at 120’000 resolution. A data-dependent acquisition method controlled by Xcalibur software (Thermo Fisher Scientific) was used that optimized the number of precursors selected (“top speed”) of charge 2+ to 5+ while maintaining a fixed scan cycle of 1.5 or 3.0 s. Peptides were fragmented by higher energy collision dissociation (HCD) with a normalized energy of 32%. The precursor isolation window used was 1.6 Th, and the MS2 scans were done in the ion trap. The m/z of fragmented precursors was then dynamically excluded from selection during 60 s. MS data were analyzed using Mascot 2.6 (Matrix Science, London, UK) set up to search the Arabidopsis thaliana Araport11 database (version of July 1st, 2015, containing 50’164 sequences, downloaded from https://araport.org), and a custom contaminant database containing the most usual environmental contaminants and enzymes used for digestion (keratins, trypsin, etc). Trypsin (cleavage at K, R) was used as the enzyme definition, allowing 2 missed cleavages. Mascot was searched with a parent ion tolerance of 10 ppm and a fragment ion mass tolerance of 0.5 Da. Carbamidomethylation of cysteine was specified in Mascot as a fixed modification. Protein N-terminal acetylation and methionine oxidation were specified as variable modifications. Scaffold (version Scaffold 4.8.4, Proteome Software Inc., Portland, OR) was used to validate MS/MS based peptide and protein identifications. Peptide identifications were accepted if they could be established at greater than 90.0 % probability by the Scaffold Local FDR algorithm. Protein identifications were accepted if they could be established at greater than 95.0 % probability and contained at least 2 identified peptides. Protein probabilities were assigned by the Protein Prophet algorithm (Nesvizhskii et al., 2003). Proteins that contained similar peptides and could not be differentiated based on MS/MS analysis alone were grouped to satisfy the principles of parsimony. Proteins sharing significant peptide evidence were grouped into clusters.

### Phylogenetic analysis

Protein multiple sequence alignments were generated with Clustal Omega (Sievers et al., 2011), and phylogenetic trees were created using the neighbor-joining method (Saitou and Nei, 1987) as implemented in SeaView (Gouy et al., 2010).

### Protein expression and purification

AtPFA-DSP1^1-215^ (UniProt, https://www.uniprot.org/ ID Q9ZVN4), AtNUDT17^23-163^ (UniProt ID Q9ZU95) and MpPFA-DSP1^4-171^ (UniProt ID A0A2R6X497) expression constructs were amplified from cDNA. A synthetic gene for MpNUDT1^18-169^ (UniProt ID A0A2R6W2U8) codon-optimized for expression in *E. coli* was obtained from Twist Bioscience. AtPFA-DSP1^1-215^ was cloned into plasmid pMH-MBP, which provides tobacco etch virus protease (TEV) cleavable N-terminal 6xHis and maltose binding protein tags. AtNUDT17^23-163^ and MpPFA-DSP1^4-171^ were cloned into pMH-HT, providing a TEV-cleavable N-terminal 6xHis tag. MpNUDT1^18-169^ was cloned into plasmid pMH-HC, providing a non-cleavable C-terminal 6xHis tag. Plasmids were transformed into *E. coli* BL21 (DE3) RIL cells. For protein expression, cells were grown in terrific broth medium at 37°C until an OD_600nm_ ∼ 0.6 and induced with 0.5 mM isopropyl-β-D-thiogalactopyranoside (IPTG) and grown at 16°C for ∼16 h. For AtPFA-DSP1 and AtNUDT17, protein expression was achieved by autoinduction. Cells were grown in terrific broth medium supplemented with lactose at 37°C until OD_600_ ∼ 0.6-0.8 and then at 16°C for 24 h. All cell pellets were harvested by centrifugation at 4,500 x g for 45 min at 4°C. AtPFA-DSP1 and AtNUDT17 were resuspended in buffer A (50 mM Tris pH 7.5, 500 mM NaCl, 1 mM DTT, Dnase I and cOmplete™ protease inhibitor cocktail [Merck]), MpPFA-DSP1 and MpNUDT1 were resuspended in buffer B (50 mM K2HPO4/KH2PO4 pH 7.8, 500 mM NaCl, 0.4 % tween, 10 mM imidazole, 10 mM β-mercaptoethanol [BME], cOmplete™ protease inhibitor cocktail [Merck]) and disrupted by sonication. Cell suspension was centrifuged at 16,000 x g for 1 h at 4°C, the supernatant was filtered through a 0.45 µm PVDF filter (Millipore) and then loaded onto an Ni^2+^ affinity column (HisTrap HP 5 mL; Cytvia) pre-equilibrated in buffer A. The column was washed with 5 column volumes of buffer A or B, respectively and fusion proteins were eluted with buffer A or B supplemented with 500 mM imidazole pH 8.0. Cleavage of the tag was performed, where applicable, by overnight incubation with TEV (1:50 ratio) at 4°C during dialysis in buffer C (20 mM Tris pH 7.5, 500 mM NaCl and 2 mM BME). The 6xHis-tagged TEV and the cleaved affinity tag were removed by a second Ni^2+^ affinity step (HisTrapExcel 5 mL; Cytvia). All samples were purified to homogeneity by size exclusion chromatography equilibrated in buffer C (20 mM Tris pH 7.5, 150 mM NaCl and 2 mM BME), on a Superdex 200 pg HR16/60 column (Cytvia) in the case of AtPFA-DSP1 and AtNUDT17, on a Superdex 200 pg HR10/30 (Cytvia) in the case of MpPFA-DSP1 and on a Superdex 75 pg HR26/60 (Cytvia) in the case of MpNUDT1. Purified proteins were snap frozen in liquid N_2_ and used for biochemical assays. Mutations were introduced by site-directed mutagenesis and the mutant proteins were purified as described for the wild type.

### Enzyme activity assays

PP-InsP phosphatase assays were performed by nuclear magnetic resonance spectroscopy (NMR). Reactions containing 100 µM of the respective [^13^C_6_]-labeled PP-InsP in 50 mM HEPES pH 7, 150 mM NaCl, 0.2 mg/mL BSA and D_2_O to a total volume of 600 µL were prepared. Reactions were supplemented with 0.5 mM MgCl_2_ were indicated. Reaction mixtures were pre-incubated at 37°C and the reaction was started by adding the respective amount of enzyme; AtPFA-DSP1 (20 nM final concentration for 1-InsP_7_, 7 nM for 5-InsP_7_, 10 nM for 1,5-InsP_8_), 1 µM of AtNUDT17, 50 nM of AtNUDT13, MpPFA-DSP1 (350 nM for 1-InsP_7_, 200 nM for 5-InsP_7_, 250 nM for 1,5-InsP_8_), 2 µM for MpPFA-DSP1^C105A^, MpNUDT1 (250 nM for 1-InsP_7_, 100 nM for 5-InsP_7_, 250 nM for 1,5-InsP_8_) or 2 µM MpNUDT1^E79A^. Reactions were monitored continuously at 37°C using a NMR pseudo-2D spin-echo difference experiment. The relative intensity changes of the C2 peaks of the respective PP-InsPs as a function of reaction time were used for quantification (Harmel et al., 2019; Zhu et al., 2019). To the raw data trend lines were added, following either a linear regression model or the first derivative of the equation of the one phase decay, normalized by the enzyme’s mass concentration.

### Western blotting

∼50 mg of leaf sample was harvested from *A. thaliana* plants and frozen into liquid N_2_ in a 2 mL Eppendorf tube with a metal bead. Samples were homogenized in a tissue lyzer (MM400, Retsch) for 30 s at a frequency of 30 Hz. Then, 50 μL of extraction buffer (100 mM Tris pH 7.5, 150 mM NaCl, 10 % [v/v] glycerol, 10 mM EDTA, 1 mM DTT, 1 mM PMSF, 1 mM Sigma protease inhibitor and 1 % [v/v] IPEGAL CA-630) were added to the tissue. Samples were mixed by vortexing, incubated for 20 min on ice and pelleted for 10 min at 20,000 x g at 4°C. The supernatant was transferred to a new tube. Protein concentrations were measured in triplicate using the Bradford protein assay in 96 well plates with 150 μL Bradford solution (Applied Chem.) and 2.5 μL of 10 times dilution of protein sample. Bovine Serum Albumine standards (0.25, 0.5, 0.75 and 1 mg/mL) were used as reference. After a 5 min incubation at room temperature (RT) in the dark, the absorbance was measured at 595 nm in a plate reader (Tecan Spark). Samples concentrations were then equalized, samples were boiled at 95°C for 5 min in SDS sample buffer, and 40 μg of protein were loaded to each lane of a 10 % SDS-PAGE tris-glycine gel. Proteins were then transferred on a nitrocellulose membrane (0.45 μm, Cytiva) for 1 h and 100 V. After blocking for 1 h with TBS-T (Tris Buffer Saline with 0.1 % Tween 20) containing 5 % (w/v) milk powder (Roth) at RT, nitrocellulose membranes were incubated at RT for 2 h with anti-GFP (Miltenyi 130-091-833) or anti-Flag (Sigma A8592) antibodies conjugated with horseradish peroxidase in TBS-T at 1:5000 dilution. Finally, after 2 washes of 5 min with TBS-T, and one wash of 5 min with TBS, blots were detected using either SuperSignal™ West Femto Maximum Sensitivity Substrate (Thermo Scientific) or BM Chemiluminescence western blotting substrate (POD; Merck) and photographic films (CL-XPosure Film, ThermoFisher). As loading control, RuBisCO was visualized using Ponceau red staining (0.1 % [w/v] Ponceau red powder (Sigma) in 5 % [v/v] acetic acid).

### Rosette size and thallus size quantification

In the case of Arabidopsis, seedling were germinated and grown on ^1/2^MS plates for 1 week and then transferred to soil for an additional 2 weeks. 15 plants per genotype (1 plant per pot) were randomized on trays. Their rosette surface areas were extracted from vertically taken images using a machine-learning approach (Ilastik, https://www.ilastik.org/) to recognize the rosette leaves. Images were segmented in Ilastik and then analyzed with Fiji (Schindelin et al., 2012).

In the case of Marchantia, thallus surface areas were quantified from single plants grown from gemmae (1 gemmae per one round 90 mm petri dish) grown in ^½^B5 medium in time course experiments defined in the respective figure legend. Image analysis was performed as described for Arabidopsis.

### PP-InsP quantification by CE-ESI-MS

Arabidopsis seedlings were grown on ^1/2^MS plates for 2 weeks and 150 mg of pooled seedling were prepared per genotype and technical replicate. Marchantia plants were grown as described above for 3 weeks and ∼500 mg of fresh weight tissue was collected for each genotype and replicate. TiO _2_ beads (Titansphere Bulk Material Titansphere 5 µm, GL Sciences; 5 mg per sample) where washed with 1 mL ddH_2_O and pelleted at 3,500 x g for 1 min at 4 °C. Beads were then washed in 1 mL of perchloric acid, pelleted again and then resuspended in 50 μL perchloric acid. Plant samples were snap frozen in liquid N_2_, homogenized by bead beating (4 mm steel beads in a tissue lyzer, MM400, Retsch), and immediately resuspended in 1 mL 1 M ice-cold perchloric acid. Samples were rotated for 15 min at 4°C and pelleted at 21,000 x g for 10 min at 4°C, the resulting supernatants were added to eppendorf tubes containing the TiO_2_ beads and mixed by vortexing. Samples were then rotated for 15 min at 4°C and pelleted at 21,000 x g for 10 min at 4°C. Beads were washed twice by resuspending in 500 μL cold 1 M perchloric acid, followed by centrifugation at 3,500 x g for 1 min at 4°C. For InsPs/PP-InsPs elution, beads were resuspended in twice 200 μL ∼2.8 % ammonium hydroxide, mixed by vortexing, rotated for 5 min and pelleted at 3,500 x g for 1 min. The two elution fractions were pooled, centrifuged at 21,000 x g for 5 min at 4°C and the supernatants were transferred to fresh tubes, and dried under vacuum evaporation for 70 min at 45-60°C. InsP/PP-InsP quantification was done utilizing an Agilent CE-QQQ system, comprising an Agilent 7100 CE, an Agilent 6495C Triple Quadrupole, and an Agilent Jet Stream electrospray ionization source, integrated with an Agilent CE-ESI-MS interface. A consistent flow rate of 10 µL/min for the sheath liquid (composed of a 50:50 mixture of isopropanol and water) was maintained using an isocratic Agilent 1200 LC pump, delivered via a splitter. Separation occurred within a fused silica capillary, 100 cm in length, with an internal diameter of 50 µm and an outside diameter of 365 µm. The background electrolyte (BGE) consisted of 40 mM ammonium acetate, adjusted to pH 9.08 with ammonium hydroxide. Before each sample run, the capillary underwent a flush with BGE for 400 seconds. Samples were injected for 15 seconds under a pressure of 100 mbar (equivalent to 30 nL). MS source parameters included a gas temperature set at 150°C, a flow rate of 11 L/min, a nebulizer pressure of 8 psi, a sheath gas temperature of 175°C, a capillary voltage of −2000V, and a nozzle voltage of 2000V. Additionally, negative high-pressure radio frequency (RF) and negative low-pressure RF were maintained at 70 and 40 V, respectively. The setting for multiple reaction monitoring (MRM) were as shown in Supplementary Figure 5B. For the preparation of the internal standard (IS) stock solution, specific concentrations were employed: 8 µM [ ^13^C_6_] 2-OH-InsP_5_, 40 µM [^13^C_6_] InsP_6_, 2 µM [^13^C_6_] 1-InsP_7_, 2 µM [^13^C_6_] 5-InsP_7_, 1 µM [^18^O_2_] 4-InsP_7_ (specifically for the assignment of 4/6-InsP_7_), and 2 µM [^13^C_6_] 1,5-InsP_8_ (Qiu et al., 2020, 2023; Haas et al., 2022). These IS compounds were introduced into the samples to facilitate isomer assignment and quantification of InsPs and PP-InsPs. Each sample was supplemented with 5 µL of the IS stock solution, thoroughly mixed with 5 µL of the sample. Quantification of InsP_8_, 5-InsP_7_, 4/6-InsP_7_, 1-InsP_7_, InsP_6_, and InsP_5_ was carried out by spiking known amounts of corresponding heavy isotopic references into the samples. Following spiking, the final concentrations within the samples were as follows: 4 μM [^13^C_6_] 2-OH-InsP_5_, 20 μM [^13^C_6_] InsP_6_, 1 μM [^13^C_6_] 5-InsP_7_, 1 μM [^13^C_6_] 1-InsP_7_, 1 μM [^18^O_2_] 4-InsP_7_, and 0.5 µM [^13^C_6_] 1,5-InsP_8_ (Harmel et al., 2019).

### β-glucuronidase (GUS) reporter assay

The β-glucuronidase (GUS) gene was used as a reporter of gene expression fused to promoters of AtPFA-DSP1, 2 and 4; AtNUDT17, 18 and 21, and MpDSP1, MpNUDT1 or MpVIP1. The previously reported _pro_AtVIH1::GUS and _pro_AtVIH2::GUS lines were used as controls (Zhu et al., 2019). 1.5-2 kbp regions upstream of the ATG were considered promoter sequences. Arabdiopsis seedling were germinated on ^½^MS medium (containing 1 % sucrose) and transferred after 1 week to ^½^MS plates containing 1% sucrose and either 0 (-Pi) or 1 mM (+Pi) K_2_HPO_4_/KH_2_PO_4_ (pH 5.7). 2-week-old seedlings were submerged in ice-cold 90 % acetone solution for 20 min and rinsed with 0.5 mM K4Fe(CN)6, 0.5 mM K3Fe(CN)6, and 50 mM NaPO4 buffer pH 7.0. Samples were then incubated in staining solution (0.5 mM K4Fe(CN)6, 0.5 mM K3Fe(CN)6, 10 mM EDTA, 0.1 % Triton X-100, 1 mM X-Gluc, and 100 mM NaPO_4_ buffer pH 7.0) and vacuum infiltrated for 15 min. Samples were placed at 37°C for the period indicated in the respective figure legend. To stop the reaction, the staining solution was replaced with aqueous solution containing increasing amounts of ethanol (15, 30, 50, 70, 100 % [v/v]) for 10 min each. Finally, the ethanol was gradually replaced by glycerol to a final concentration of 30 % (v/v) before recording images in a binocular (Nikon SMZ18 equipped with a DS-Fi3 CMOS camera). In the case of Marchantia, plants from single gemmae were grown for 1 week on ^1/2^B5 medium and transferred to plates containing either 0 (-Pi) or 0.5 mM (+Pi) K_2_HPO_4_/KH_2_PO_4_ (pH 5.5). The same staining protocol was used as described for Arabidopsis.

### RNA-seq analyses

2-week-old Arabdiopsis and Marchantia plants grown under Pi-sufficient or Pi-starvation conditions (as described in the β-glucuronidase (GUS) reporter assay section). For each biological replicate, 3-4 plants were pooled and RNA was extracted using the RNeasy plant mini kit (Qiagen). 100 ng of total RNA per sample determined using a Qubit fluorometer (Thermofisher). RNA quality control using 2100 Bioanalyzer system (Agilent Technologies), library preparation and sequencing were performed by the *iGE3* Genomic Platform at the Faculty of Medicine, University of Geneva (https://ige3.genomics.unige.ch/). Sequencing was performed with Novaseq 6000 machine from Illumina with 100 bp single-read output. Quality control of the reads and adaptor trimming were done with MultiQC (Ewels et al., 2016). Genomic and transcript annotation files of the *Arabidopsis thaliana* TAIR10 reference genome were downloaded from the TAIR database (https://www.arabidopsis.org/). In the case of *Marchantia polymorpha*, the v6.1 reference genome and annotation were downloaded from MarpolBase (https://marchantia.info/download/MpTak_v6.1/). For mapping the reads, HISAT2 (Kim et al., 2019) (v2.2.1 with only the -dta option in extra) and StringTie (Pertea et al., 2016) (v2.2.1 with default options) were used. Ballgown (Pertea et al., 2016) was used to re-assemble the different output files into a single tab-delimited file. Prior to further statistical analysis, counts were filtered to have at least 10 counts per gene in at least one sample. DESeq2 (Love et al., 2014) (v3.17) with default options has been used in Rstudio (https://posit.co/download/rstudio-desktop/) to make pairwise comparison of the different genotype and growth conditions vs. the Col-0 (Arabidopsis) or Tak-1 (Marchantia) references, respectively. Gene ontology enrichment analyses were performed in Panther (https://www.pantherdb.org/), data visualization was done in R (R Core Team, 2014) packages ggplot2, dplyr, reshape2 and EnhancedVolcano. Raw reads have been deposited with the sequence read archive (SRA; https://submit.ncbi.nlm.nih.gov/subs/sra/), with identifiers PRJNA1090032, PRJNA1088982, PRJNA1089142 and PRJNA1090651.

### Phosphate and nitrate quantification

Arabidopsis seedlings were germinated on ^½^MS supplemented with 1 % (w/v) sucrose for one week and then transferred to ^½^MS agar plates supplemented with 1 % (w/v) sucrose, containing either 0 mM Pi (-Pi), 1 mM KH_2_PO_4_/K_2_HPO_4_ (pH 5.7) or 2 mM Pi (+Pi). At 2 weeks, four seedlings were pooled, weighed, resuspended in 400 µL miliQ H_2_O in an Eppendorf tube and snap-frozen in liquid N_2_. Plants were homogenized using a tissue lyzer (MM400, Retsch) and then samples were thawed at 85°C for 15 min with orbital shaking and snap frozen again in liquid N_2_. Samples were thawed again at 85°C for 1 h with orbital shaking. Free inorganic phosphate concentrations were determined by a colorimetric molybdate assay (Ames, 1966). The master mix for each sample contained 72 µL of ammonium molybdate solution (0.0044 % [w/v] of ammonium molybdate tetra hydrate, 0.23 % [v/v] of 18 M H_2_SO_4_), 16 µL of 10 % (w/v) acetic acid and 12 µL of miliQ H_2_O. For each sample, 100 µL of the mix was incubated with 20 µL of each sample in a 96-wells plate. Standard curves obtained by diluting 100 mM Na_2_HPO_4_ solution to final concentrations of 2, 1, 0.5, 0.25, 0.16 and 0.08 mM. Technical triplicates were done for the standards and duplicates for all samples. Plates were incubated for 1 h at 37°C and absorbance at 820 nm was measured using a Spark plate reader (Tecan).

In the case of Marchantia, plants were grown from gemmae for 1 week on ^½^B5 medium. Plants were then transferred to plates containing either 0 mM Pi (-Pi) or 0.5 mM KH_2_PO_4_/K_2_HPO4 (pH 5.5) (+Pi). One plant represents one biological replicate, samples were processed as described for Arabidopsis above.

Nitrate quantification were based on the Miranda colorimetric assay (Miranda et al., 2001). Marchantia plants were grown on ^½^B5 medium and processed as described above. Miranda solution (0.25 % [w/v] vanadium III chloride, 0.1 % [w/v] sulfanilamide and 0.1 % [w/v] N-(1-naphthyl)ethylenediamine in 0.5 M HCl) was prepared and 200 µL of the solution was mixed with 5 µL for each sample in a 96-wells plate. Standards were prepared by diluting KNO_3_ to final concentrations of 1, 0.5, 0.25, 0.12 and 0.06 mM. Technical triplicates were done for standards and duplicate for all samples. Plates were incubated at 65°C for 2 h and the absorbance at 540 nm was measured using a Spark plate reader (Tecan).

### Elemental quantifications

Plants were grown from gemmae as described above for 3 weeks on ^½^B5 medium. ∼ 8 g of plant material was harvested for each genotype, rinsed in aqueous solution containing 10 mM EDTA for 10 min with gentle shaking. Samples were rinsed 3 times with milQ H_2_O for 5 min and dried at 65°C for 2 d. For the different ion quantifications samples were then split into 20 mg batches. Each batch was incubated overnight with 750 μL of nitric acid (65 % [v/v]) and 250 μL of hydrogen peroxide (30 % [v/v]). Next, samples were mineralized at 85°C for 24 h. Finally, milliQ H_2_O was added to each sample and the elemental quantifications were done using inductively coupled plasma optical emission spectrometer (ICP-OES 5800, Agilent Technologies).

### Marchantia histology

The portion of interest of the plant was sectioned and fixed in phosphate buffer pH 7.2 with 4 % (w/v) formaldehyde, 0.25 % (w/v) glutaraldehyde and 0.2 % (v/v) Triton X-100; the fixation was done overnight at 4°C under agitation, after vacuum infiltration. Samples were then washed with phosphate buffer (2×15 min) and with water (2×10 min) before undergoing dehydration in a graded ethanol series (ethanol 30 %, 50 %, 70 %, 90 % and 100 % with incubations respectively of 30 min, 2×30 min, 3×20 min, 2×30 min, 2×30 min and overnight at 4°C in the last bath of ethanol 100 %). Technovit 7100 was prepared according to the manufacturer’s indications by supplementing it with Hardener I (product from the kit), and samples were progressively infiltrated by incubations in 3:1, 1:1 and 1:3 mixes ethanol:Technovit 7100 (each time 2 hours under agitation at room temperature), before finally incubating in 100 % Technovit 7100 for 2 hours at room temperature (after vacuum infiltration) and for another 40 hours at 4°C. Embedding was done in Technovit 7100 supplemented with 1/15 Hardener II and 1/25 polyethylene glycol 400; polymerization was done for 30 min at room temperature followed by 30 min at 60°C. Sectioning was performed with a Histocore AUTOCUT microtome (Leica) using disposable R35 blades and sections of 4 µm were deposited on SuperFrost slides.

For the ruthenium red staining, sections were incubated 1 minute in 0.05 % (w/v) ruthenium red in distilled water, extensively rinsed with distilled water, then incubated 1 minute in xylene and mounted in Pertex. For the fluorol yellow staining, sections were incubated 5 min in 0.01 % (w/v) fluorol yellow in 50 % ethanol, extensively rinsed with distilled water, washed with distilled water during 1 hour with agitation and mounted in 50 % glycerol. For the Renaissance SR2200 staining, sections were incubated 1 minute in 0.05 % (v/v) Renaissance SR2200, extensively rinsed with distilled water, washed with distilled water during 1 hour with agitation and mounted in phosphate buffer saline pH 7.4 supplemented with 50 % glycerol.

Sections stained with ruthenium red were observed with a Leica DM6B widefield microscope equipped with a DMC5400 CMOS camera (used with binning 2×2) and a 20x Fluotar NA 0.55 air objective. Sections stained with fluorol yellow and Renaissance SR2200 were observed with a Leica TCS SP8 confocal system mounted on a DMi8 inverted microscope and in the following configuration: objective HC PL APO CS2 20x NA 0.75 IMM used with water immersion; sampling speed 400 Hz; pixel size 190 nm; pinhole 1.0 Airy Unit (fluorol yellow) or 1.5 Airy Unit (Renaissance SR2200); frame averaging 3. Fluorol yellow and Renaissance SR 2200 were excited at 488 nm and 405 nm respectively, and their fluorescence was collected by a HyD detector at gain 50 % between 507 nm and 550 nm (fluorol yellow) and between 420 nm and 480 nm (Renaissance SR2200). For a given dye, all images were acquired and post-processed identically.

Image processing was done using Fiji (Schindelin et al., 2012). For ruthenium red pictures, tiling and stitching was done using the Leica LAS X Navigator tool. For fluorol yellow pictures, a Gaussian blur of radius 1 pixel was applied, the images were downscaled using bicubic interpolation (from 4096×4096 to 1024×1024) and finally a rolling ball background subtraction was applied with a radius of 50 pixels; the look-up table NanoJ-Orange was used to display the images. For Renaissance SR2200 pictures, a Gaussian blur of radius 0.6 pixel was applied and the images were downscaled using bicubic interpolation; the look-up table Fire was used.

## Acknowledgments

We thank D. Furkert for providing the 5PCP-InsP_5_ resin, J. Santiago, M. Barberon and P. Rieu for critical reading of the manuscript, B. Petit and M. Docquier from the *iGE3* Genomic Platform at the Faculty of Medicine, University of Geneva for performing the different RNA-seq experiments, and the Service d’Analyses Multi-Elementaires (SAME) from the University of Montpellier, INRAE, CNRS, Institut Agro, Montpellier, France for elemental quantifications. This work was supported by the European Union ERC consolidator grant 818696 INSPIRE (to M.H.), by Swiss National Science Foundation Sinergia grant CRSII5_209412 (to M.H. and D.F.), by an International Research Scholar Award by the Howard Hughes Medical Institute (to M.H.), and by the Deutsche Forschungsgemeinschaft (DFG) under Germany’s Excellence Strategy CIBSS, EXC-2189, Project ID 390939984 (to H.J.).

## Author contributions

M.H. designed the project, with input from D.C.; F.L. and M.H. conceived experiments; F.L. expressed and purified proteins, performed western blots, reporter GUS assays, RNA-seq experiments, generated and analyzed transgenic lines in Arabidopsis, with help from J. N., and generated and analyzed Marchantia transgenics with help from F.R. and J.N.; S.B. performed NMR-based enzyme assays; A.S. performed PP-InsP quantifications with samples from F.L.; D.C. performed the PP-InsP interaction screen; C.F. performed Marchantia histology; F.L. quantified plant phenotypes, phosphate and nitrate levels, and analyzed RNA-seq data; S.L. analyzed histological data; D.F. and H.J. provided PP-InsP reagents and analyzed NMR and CE-ESI-MS data. M.H. and F.L. wrote the paper, with input from S.B., A.S., F.R., S.L., H.J. and D.F..

---

**Supplementary Figure 1.**
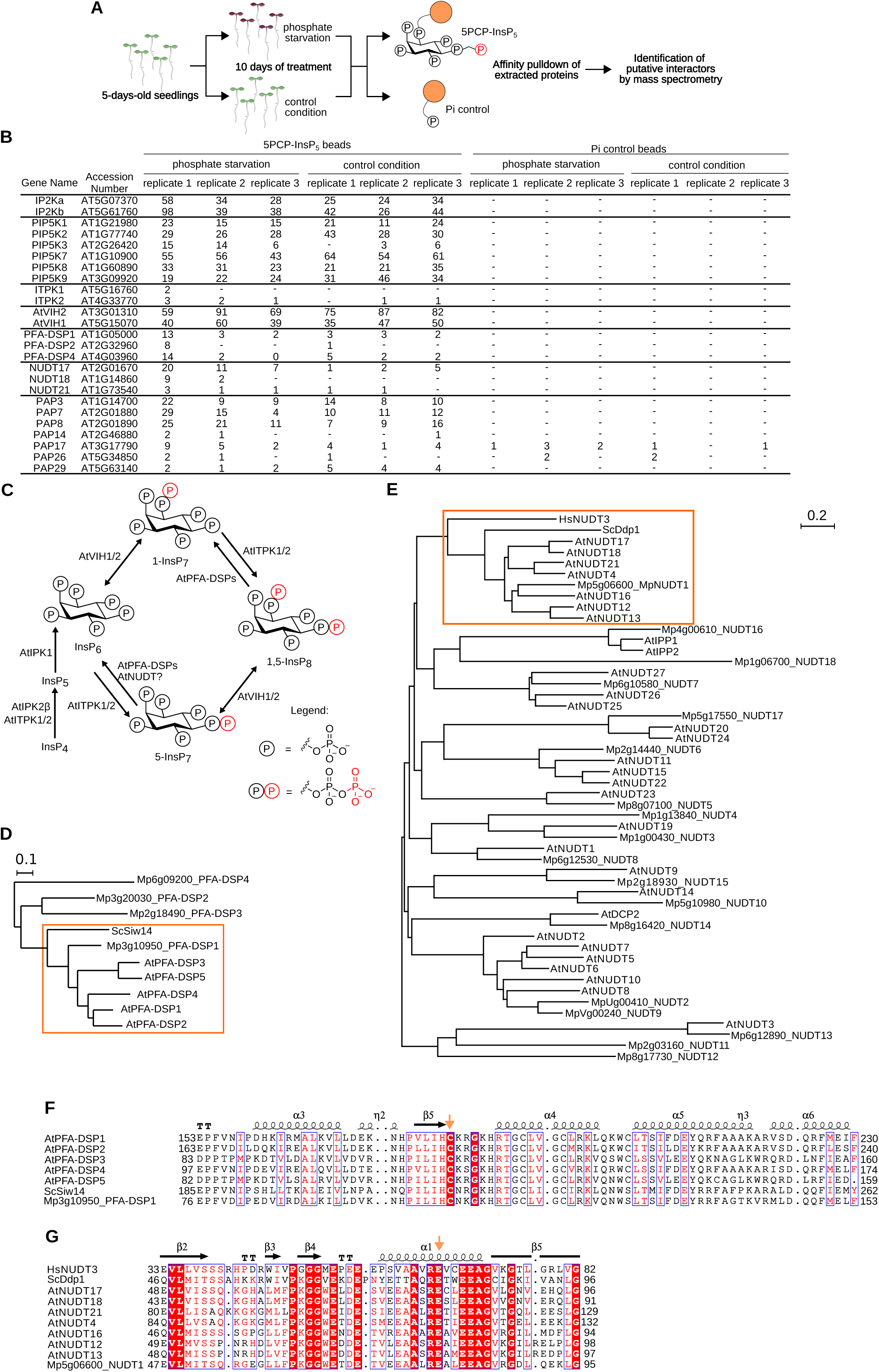
5PCP-InsP_5_ interaction screen identifies putative PP-InsPs pyrophosphate phosphatases in Arabidopsis, related to Figure 1. **(A)** Schematic overview of the interaction screen. Col-0 seedlings were germinated on ^½^MS plates for 5 d, and then transferred to liquid ^½^MS medium (containing 1 % [w/v] sucrose) in the presence of 0.2 µM (-Pi) or 1 mM (+Pi) K_2_HPO_4_/KH_2_PO_4_ (pH 5.7) for 10 d. **(B)** Table summary of all known and putative PP-InsP kinases and phosphatases recovered from the 5PCP-InsP_5_ screen described in **(A)**. Peptide counts are shown alongside. **(C)** Schematic overview of the PP-InsP biosynthesis and catabolic pathway in Arabidopsis. **(D-E)** Phylogenetic trees of PFA-DSPs (AtPFA-DSP1 UniProt, https://www.uniprot.org/ ID Q9ZVN4, AtPFA-DSP2 Q84MD6, AtPFA-DSP3 Q681Z2, AtPFA-DSP4 Q940L5, AtPFA-DSP5 Q9FFD7, ScSiw14 P53965, MpPFA-DSP accession numbers from http://marchantia.info) **(D)** or NUDT (AtNUDT4 Q9LE73, AtNUDT12 Q93ZY7, AtNUDT13 Q52K88, AtNUDT16 Q9LHK1, AtNUDT17 Q9ZU95, AtNUDT18 Q9LQU5, AtNUDT21 Q8VY81, ScDdp1 Q99321, HsNUDT3 O95989) **(E)** enzymes present in *A. thaliana*, *M. polymorpha, S. cerevisiae* or *H. sapiens*. Subtrees containing the respective enzymes identified in the 5PCP-InsP_5_ screen are marked with a orange rectangle. **(F-G)** Multiple sequence alignment of the selected PFA-DSPs **(F)** or NUDT **(G)** enzyme family members. The crystal structure of AtPFA-DSP1 (http://rcsg.org PDB-ID: 1XRI) or HsNUDT3 (PDB-ID: 2FVV) were used to generate the secondary structure assignments. Catalytic residues targeted by site-directed mutagenesis in Figure 3 are marked by an arrow (shown in orange).

**Supplementary Figure 2.**
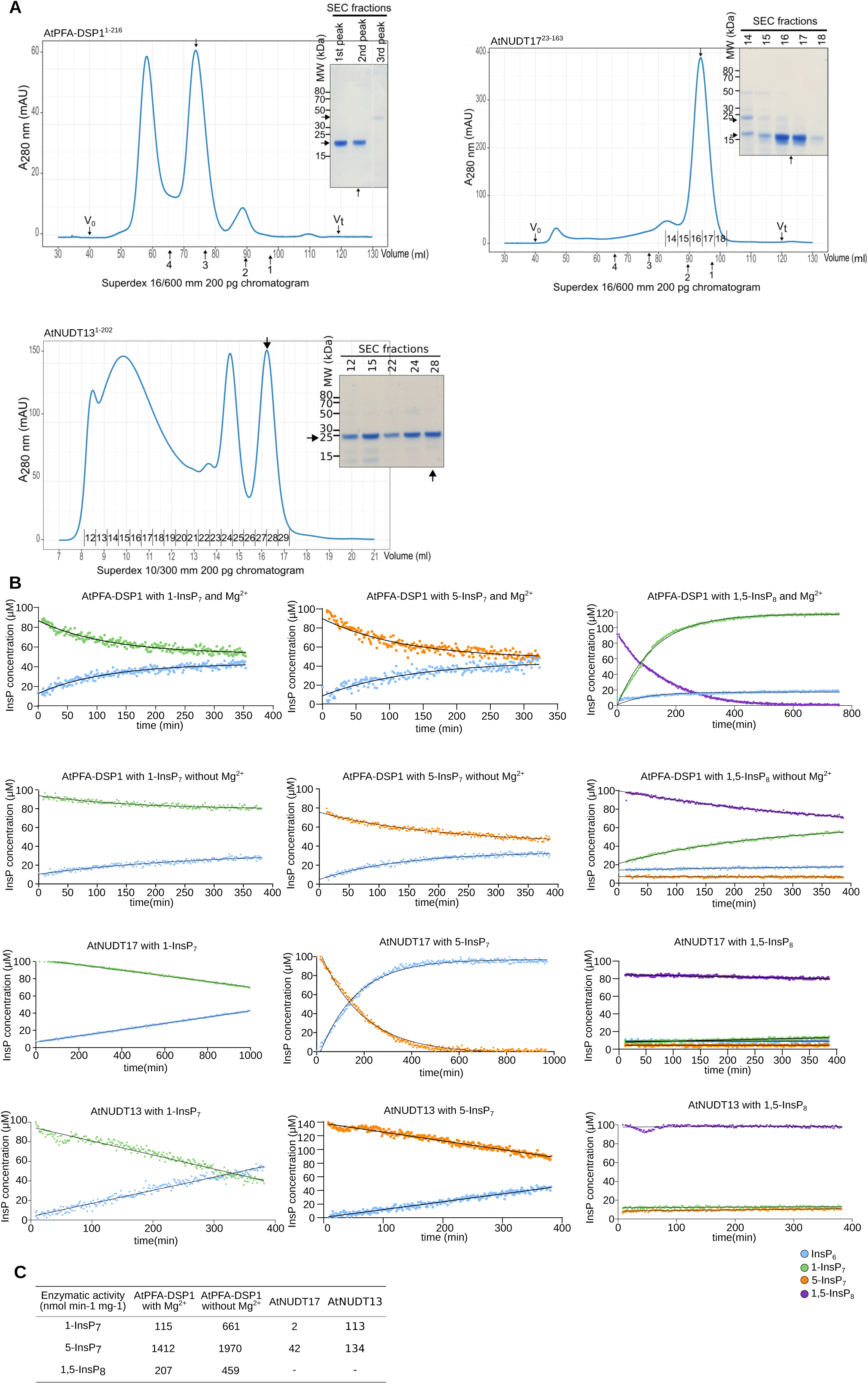
Purification and inositol pyrophosphate phosphatase activities of recombinant AtPFA-DSP1, AtNUDT17 and AtNUDT13, related to Figure 1. **(A)** Size exclusion chromatography chromatograms of purified AtPFA-DSP1^1-216^ AtNUDT17^23-163^ and AtNUDT13^1-202^. Arrows indicate the elution volumes of protein standards: 1: ribonuclease A (13.7 kDa), 2: carbonic anhydrase (29 kDa), 3: conalbumin (75 kDa) and 4: ferritin (440 kDa). The calculated theoretical molecular masses are: AtPFA-DSP1^1-216^ ∼24 kDa, AtNUDT17^23-163^ ∼16 kDa, AtNUDT13^1-202^ ∼24 kDa, MBP ∼45 kDa and TEV ∼25 kDa. Coomassie-stained SDS-PAGE analyses of the peak fractions are shown alongside. **(B)** NMR time course experiments of AtPFA-DSP1, AtNUDT17 and AtNUDT13 using 100 μM of [^13^C_6_]-labeled PP-InsP as substrate. Reactions had a different amount of protein depending on the couple protein/substrate used. Pseudo-2D spin-echo difference experiments were used and changes in the relative intensities of the C2 peaks of the respective InsPs were quantified. **(C)** Table summaries of the enzyme activities for AtPFA-DSP1 (either in the presence or absence of 0.5 mM MgCl_2_), AtNUDT17 and AtNUDT13.

**Supplementary Figure 3.**
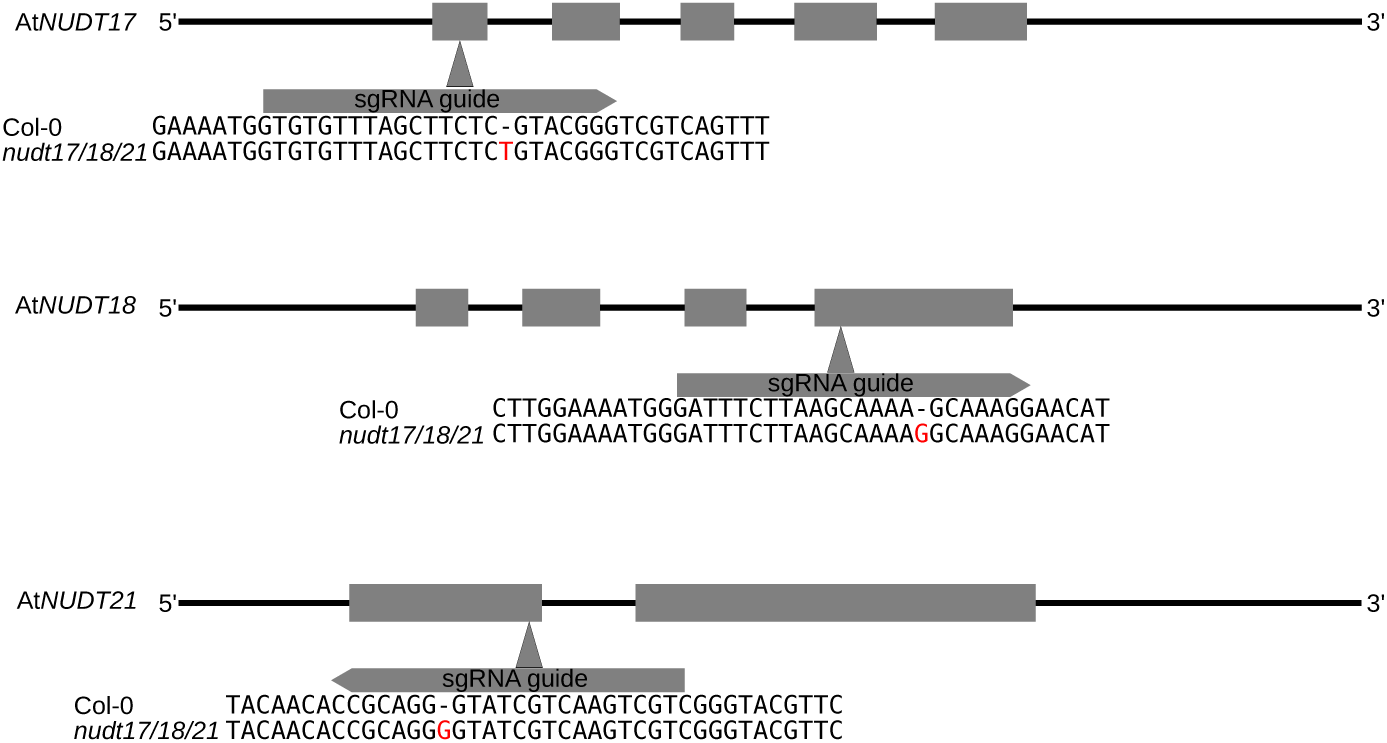
CRISPR/Cas9 gene editing events in the *nudt17/18/21* mutant, related to Figure 1. Schematic overview of the At*NUDT17*, At*NUDT18* and At*NUDT21* genes with exons depicted as squares and introns as lines. CRISPR-Cas9 sgRNA guide sequences are shown alongside, all causing single base insertion events, as confirmed by Sanger sequencing.

**Supplementary Figure 4.**
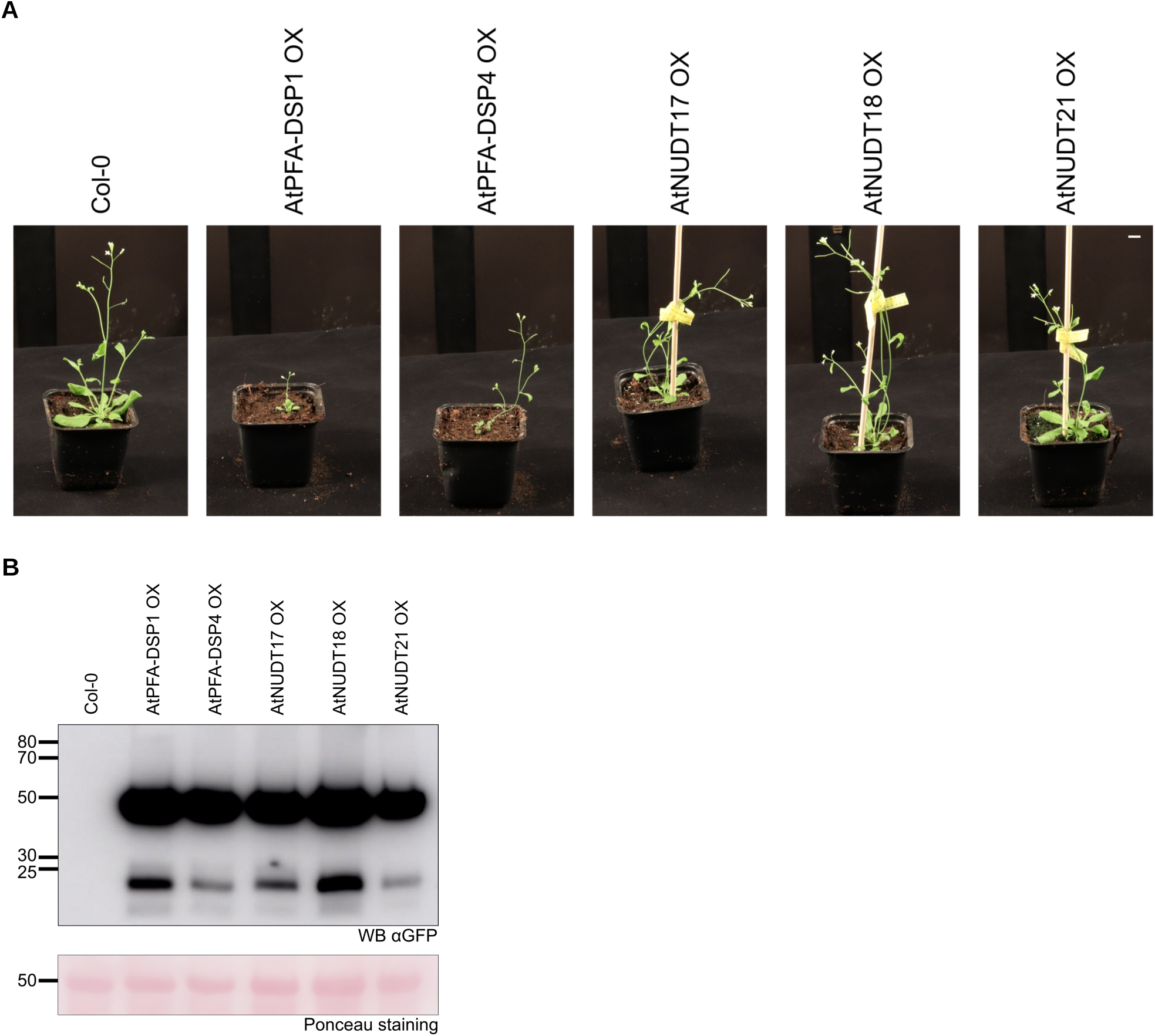
Growth phenotypes of At*PFA-DSP1* OX, At*PFA-DSP4* OX, At*NUDT17 OX*, At*NUD18* OX and At*NUDT21* OX lines, related to Figure 1. **(A)** Growth phenotypes of 4-week-old At*PFA-DSP1* OX, At*PFA-DSP4* OX, At*NUDT17 OX*, At*NUD18* OX and At*NUDT21* OX plants, all expressed from the constitutive Ubiquitin 10 promoter and carrying a C-terminal GFP tag. Plants were germinated on ^½^MS for 1 week before transfer to soil (scale bar = 1 cm). **(B)** Western blot of the plants described in **(A)** with a ponceau stain shown below as loading control.

**Supplementary Figure 5.**
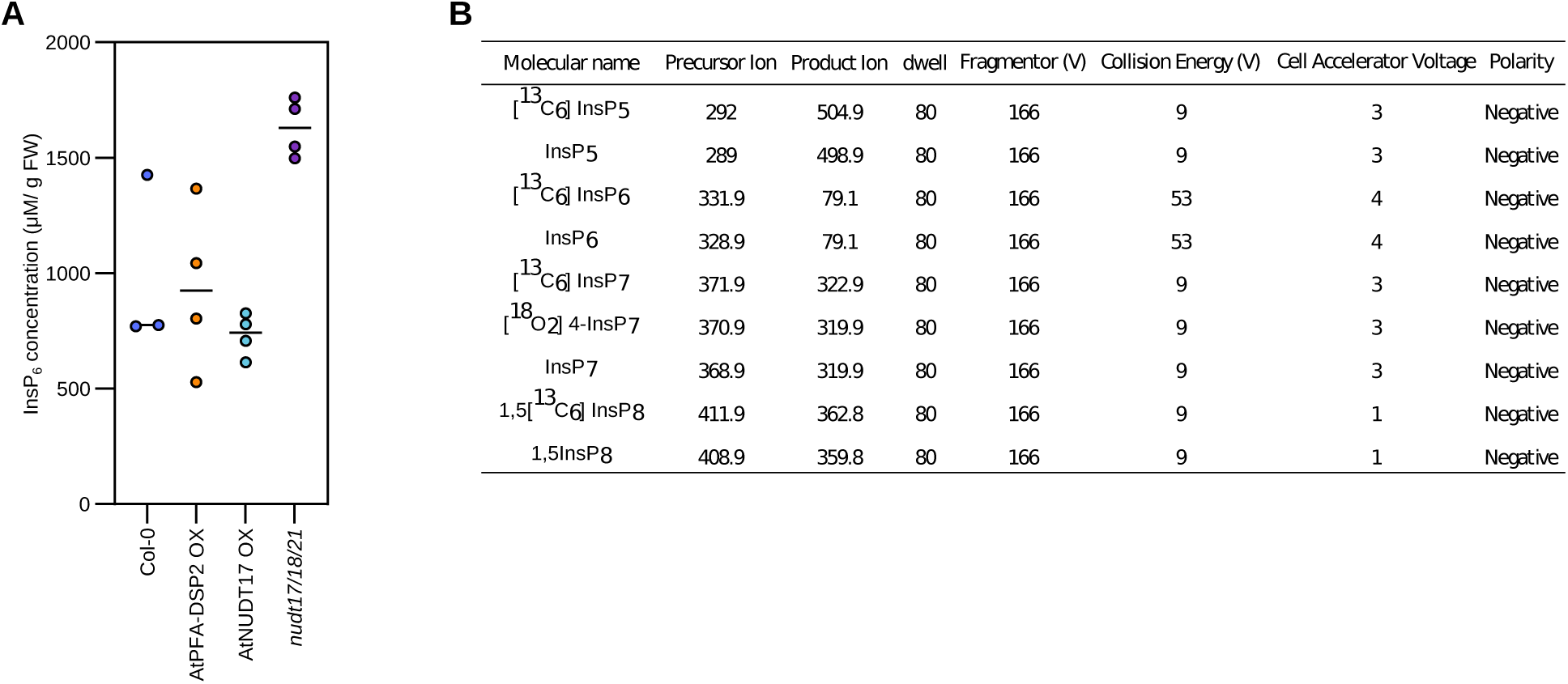
InsP_6_ levels in wild-type and transgenic Arabidopsis plants, related to Figure 1. **(A)** InsP_6_ concentrations for Col-0, *nudt17/18/21*, At*PFA-DSP2* OX and At*NUDT17* OX plants were determined using the CE-ESI-MS method and seedlings grown on ^½^MS for 2 weeks. InsP_6_ levels were normalized by fresh weight. **(B)** Mass spectrometry parameters table for multiple reaction monitoring transitions.

**Supplementary Figure 6.**
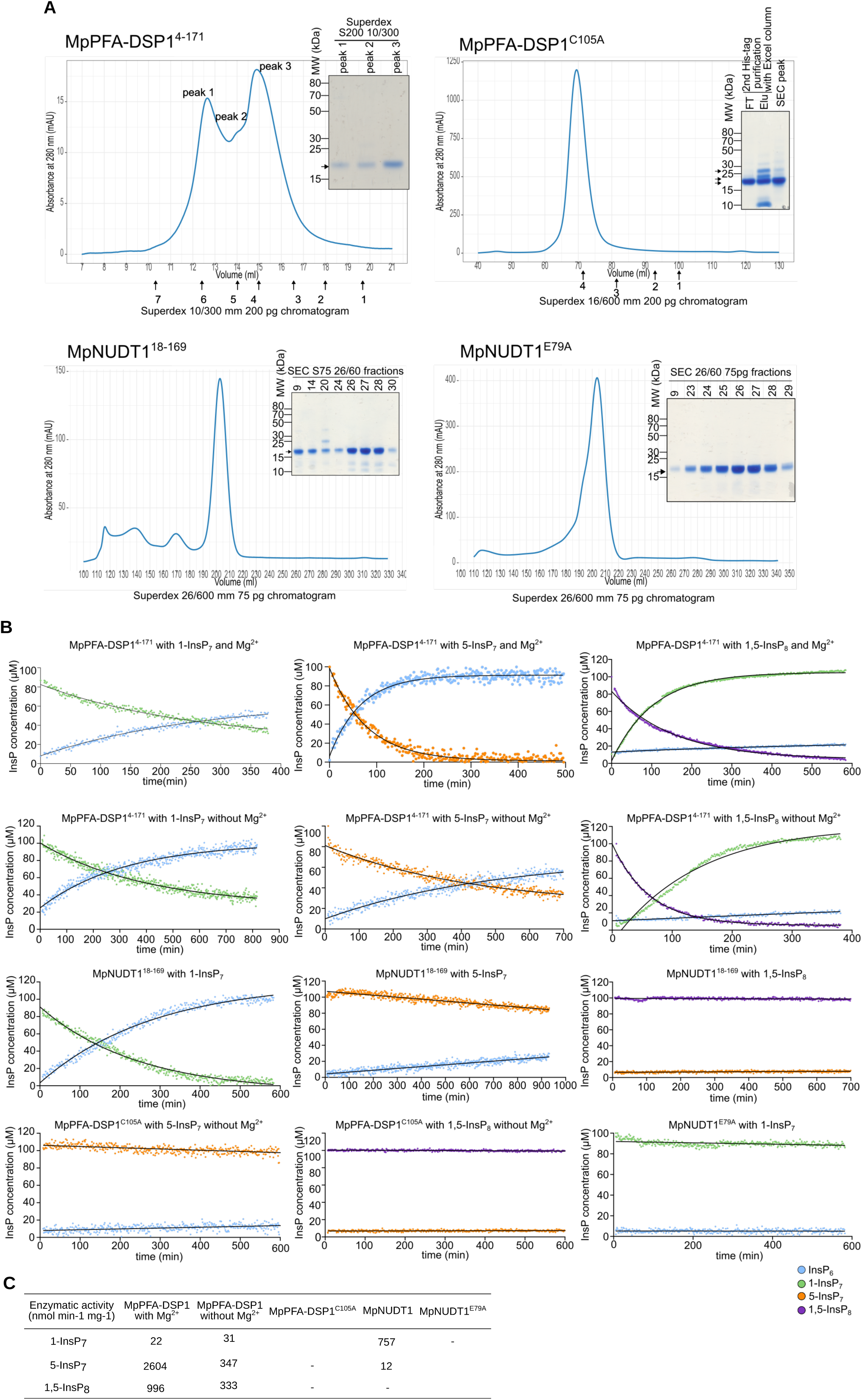
Purification and inositol pyrophosphate phosphatase activities of recombinant MpPFA-DSP1 and MpNUDT1, related to Figure 3. **(A)** Size exclusion chromatography traces of purified MpPFA-DSP1^4-171^, MpPFA-DSP1^C105A^, MpNUDT1^18-169^ and MpNUDT1^E79A^. Arrows indicate the elution volume of standards: 1: aprotinin (6.5 kDa), 2: ribonuclease A (13.7 kDa), 3: carbonic anhydrase (29 kDa), 4: ovalbumin (44 kDa), 5: conalbumin (75 kDa), 6: aldolase (158 kDa) and 7: ferritin (440 kDa). The calculated theoretical molecular masses are: MpPFA-DSP1^4-171^ ∼20 kDa, HT-MpPFA-DSP1^4-171^ ∼23 kDa and HC-MpNUDT1^18-169^ ∼19 kDa. Coomassie-stained SDS PAGE analyses of the peak fractions are shown alongside. **(B)** NMR time course experiments of MpPFA-DSP1^4-171^, MpPFA-DSP1^C105A^, MpNUDT1^18-169^ and MpNUDT1^E79A^ using 100 μM of [^13^C_6_]-labeled PP-InsP as substrate. **(C)** Table summaries of the enzyme activities for MpPFA-DSP1^4-171^, MpPFA-DSP1^C105A^, MpNUDT1^18-169^ and MpNUDT1^E79A^ toward the different PP-InsP isomers.

**Supplementary Figure 7.**
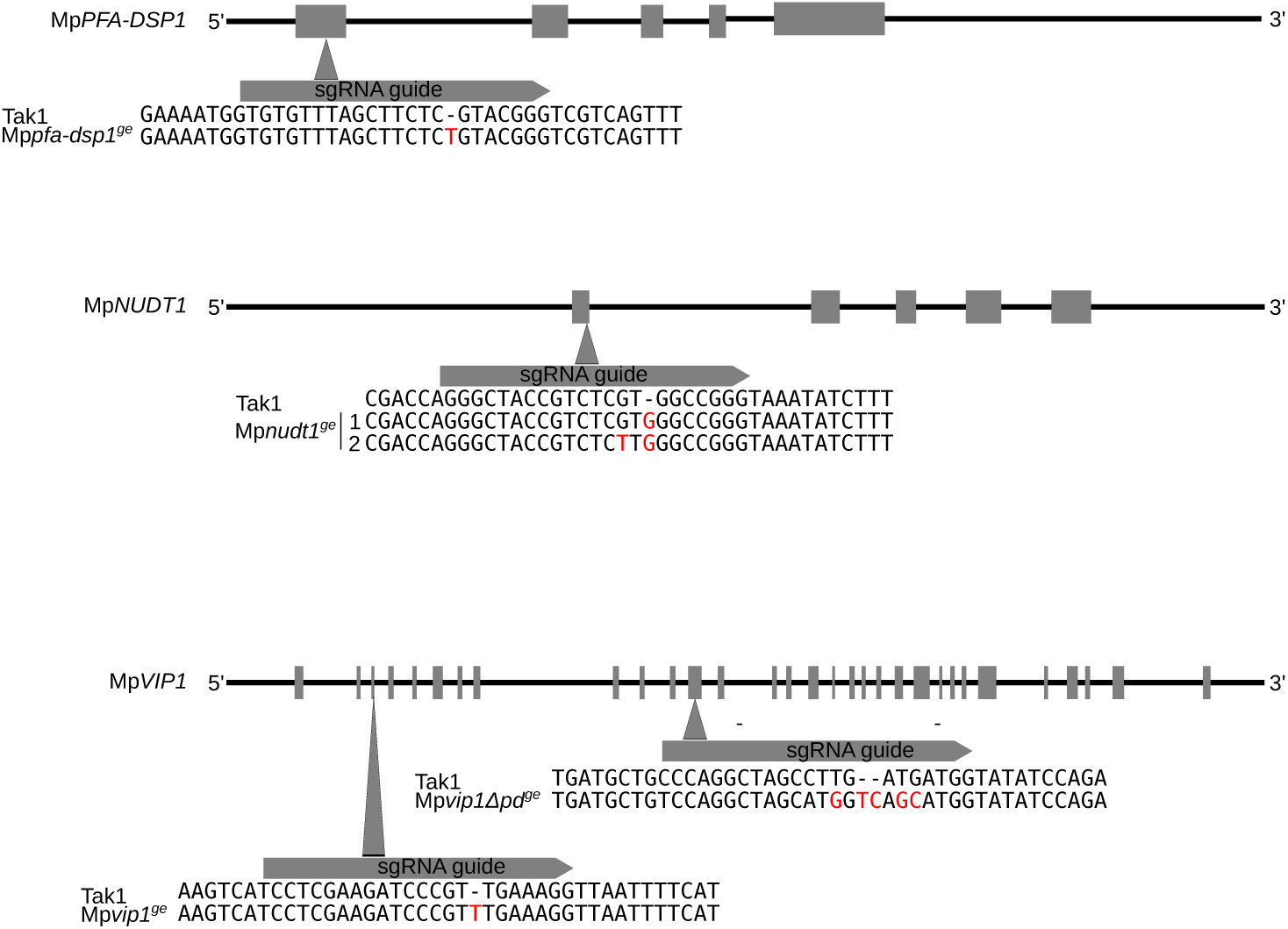
CRISPR/Cas9 gene editing events in the Mp*pfa-dsp1^ge^*, Mp*nudt1^ge^*, Mp*vip1^ge^*, Mp*vipΔpd^ge^* mutants, related to Figure 3. Schematic overview of Mp*PFA-DSP1*, Mp*NUDT1* and Mp*VIP1* genes with the exons depicted as squares and introns and UTRs as lines. CRISPR-Cas9 sgRNA guide sequences are shown alongside, all causing single base insertion events, as confirmed by Sanger sequencing.

**Supplementary Figure 8.**
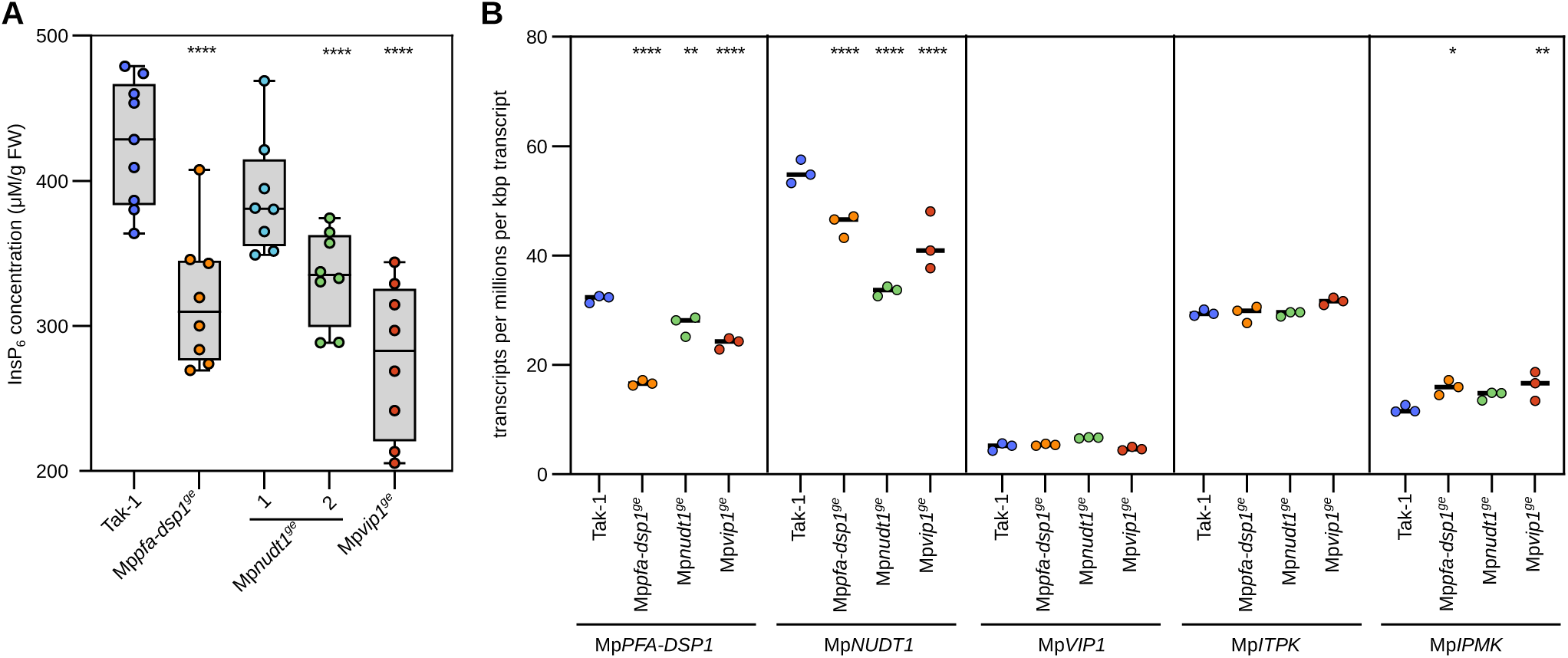
InsP_6_ and PP-InsP levels in wild-type and transgenic Marchantia plants, related to Figure 3. **(A)** InsP_6_ concentrations for Tak-1, Mp*pfa-dsp1^ge^*, Mp*nudt1^ge^*, Mp*vip1^ge^* were determined using the CE-ESI-MS method and plants grown on ^½^B5 for 3 weeks. InsP_6_ levels were normalized by fresh weight. **(B)** RNA-seq derived gene expression of Mp*PFA-DSP1*, Mp*NUDT1*, Mp*VIP1*, Mp*ITPK* (the putative InsP_6_ kinase in *M. polymorpha*) and Mp*IPMK*, comparing 3-week-old Mp*pfa-dsp1^ge^*, Mp*nudt1^ge^* and Mp*vip1^ge^* plants grown under Pi-sufficient conditions to the Tak-1 wild type.

**Supplementary Figure 9.**
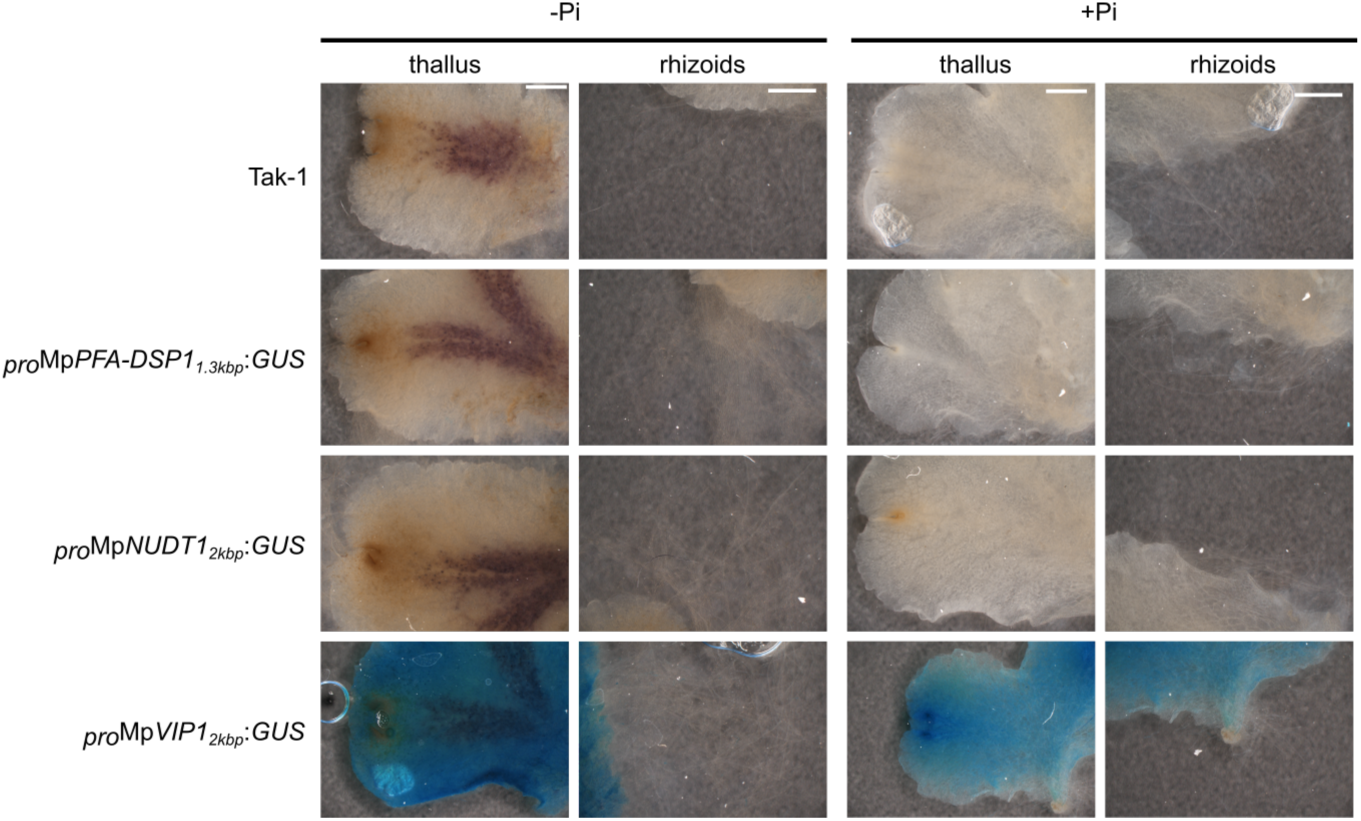
β-glucuronidase (GUS) assay for different Marchantia reporter lines grown under Pi-sufficient or Pi-starvation conditions Pi starvation, related to Figure 4. Transgenic lines expressing β-glucuronidase (GUS) gene fused to the promoters of Mp*PFA-DSP1*, Mp*NUDT1* and Mp*VIP1* were grown from gemmae for one week on ^½^B5 medium plates and then transferred ^½^B5 medium plates containing either 0 mM (-Pi) or 0.5 mM K_2_HPO_4_/KH_2_PO_4_ (pH 5.7) (+Pi) for another week. Samples were stained for 4 h and analyzed for β-glucuronidase activity (scale bar = 0.1 cm).

**Supplementary Figure 10.**
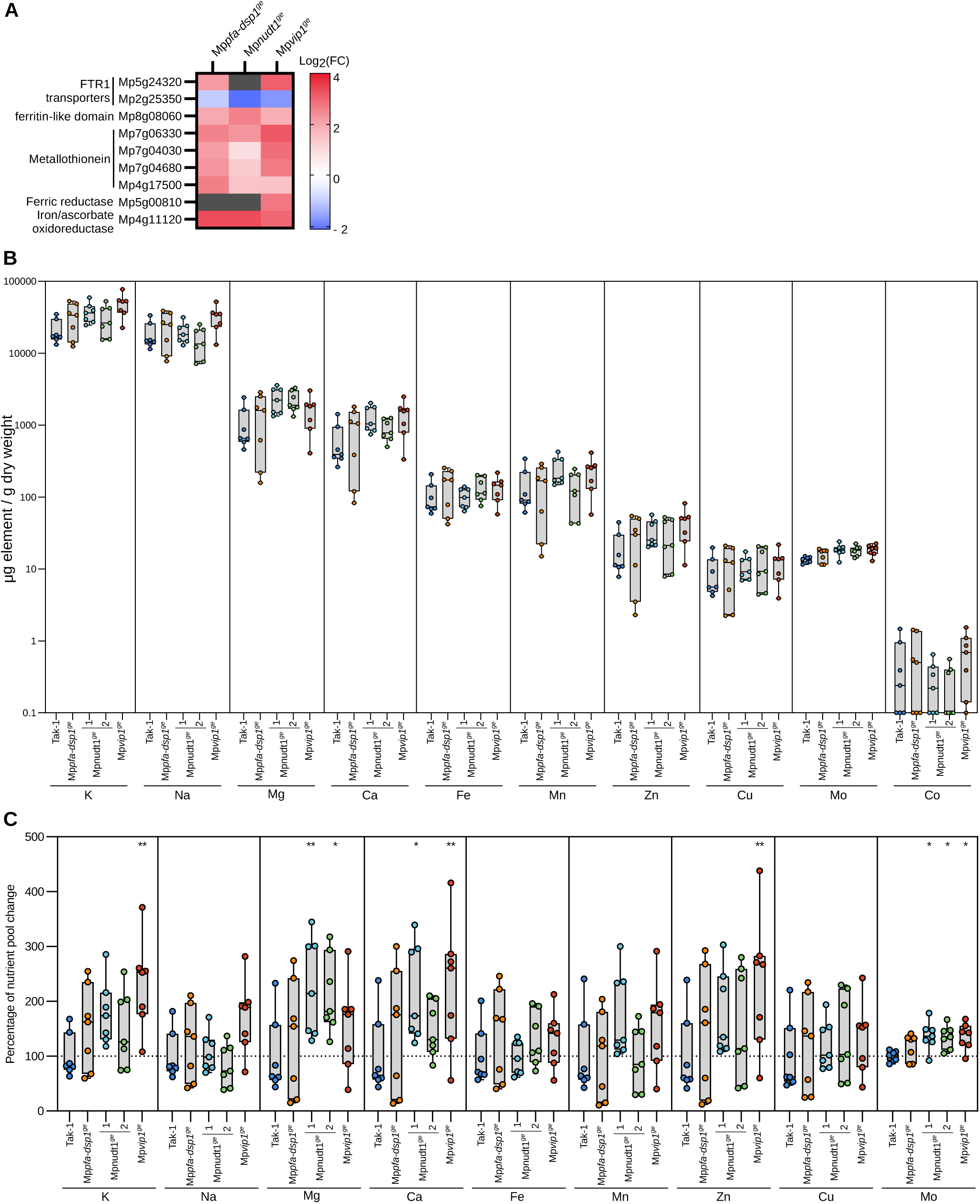
Metal ion homeostasis is not severely affected in Mp*pfa-dsp1^ge^*, Mp*nudt1^ge^* or Mp*vip1^ge^* mutants, related to Figure 4. **(A)** Heatmap of differentially expressed genes (DEGs) in 3-week-old Mp*pfa-dsp1^ge^*, Mp*nudt1^ge^* and Mp*vip1^ge^* mutant plants vs. Tak-1 grown under Pi-sufficient conditions. Reads were mapped to the reference genome with HISAT2 and and DEGs were obtained with DESeq2 with a filter limit of a minimum of 10 reads per dataset. Genes significantly different from Tak-1 involved in metal ions homeostasis are displayed. Grey boxes = no differential expression. **(B-C)** Ionomic profiles of Tak-1, Mp*pfa-dsp1^ge^*, Mp*nudt1^ge^* and Mp*vip1^ge^* plants. Plants were grown from gemmae for 3 weeks on ^½^B5 medium plates. Each replicate had ∼20 mg of dry weight. Ionomic profiling was performed by inductively coupled plasma optical emission spectrometer (ICP-OES 5800, Agilent Technologies) with 3 technical replicates per biological sample. Is shown first the raw data of μg of element per g of dry weight in **(B)** and normalized by Tak-1 average for each element in **(C)**. A Dunnett (Dunnett, 1955) test was performed for each element with Tak-1 as reference in **(C)**.

